# A Human-Immune-System (HIS) humanized mouse model (DRAGA: HLA-A2. HLA-DR4. Rag1 KO.IL-2Rγc KO. NOD) for COVID-19

**DOI:** 10.1101/2020.08.19.251249

**Authors:** Teodor-D. Brumeanu, Pooja Vir, Ahmad Faisal Karim, Swagata Kar, Dalia Benetiene, Megan Lok, Jack Greenhouse, Tammy Putmon-Taylor, Christopher Kitajewski, Kevin K. Chung, Kathleen P. Pratt, Sofia A. Casares

**Affiliations:** Uniformed Services University of the Health Sciences, Department of Medicine, Division of Immunology, Bethesda, MD 20814, U.S.A.; Naval Medical Research Center/Walter Reed Army Institute of Research, Infectious Diseases Directorate, Silver Spring, MD 20910, U.S.A.; Bioqual Inc., Rockville, MD 20852, U.S.A.

**Keywords:** SARS-CoV-2, HIS-humanized mouse model, COVID-19, lung immunopathology

## Abstract

We report the first Human Immune System (HIS)-humanized mouse model (“DRAGA”: HLA-A2.HLA-DR4.Rag1KO.IL-2RγcKO.NOD) for COVID-19 research. This mouse is reconstituted with human cord blood-derived, HLA-matched hematopoietic stem cells. It engrafts human epi/endothelial cells expressing the human ACE2 receptor for SARS-CoV-2 and TMPRSS2 serine protease co-localized on lung epithelia. HIS-DRAGA mice sustained SARS-CoV-2 infection, showing deteriorated clinical condition, replicating virus in the lungs, and human-like lung immunopathology including T-cell infiltrates, microthrombi and pulmonary sequelae. Among T-cell infiltrates, lung-resident (CD103^+^) CD8^+^ T cells were sequestered in epithelial (CD326^+^) lung niches and secreted granzyme B and perforin, indicating cytotoxic potential. Infected mice also developed antibodies against the SARS-CoV-2 viral proteins. Hence, HIS-DRAGA mice showed unique advantages as a surrogate *in vivo* human model for studying SARS-CoV-2 immunopathology and for testing the safety and efficacy of candidate vaccines and therapeutics.

---

Infection with Severe Acute Respiratory Syndrome-coronavirus-2 (SARS-CoV-2), the highly transmissible pathogen responsible for the ongoing pandemic coronavirus disease 2019 (COVID-19), results in outcomes from asymptomatic or mild disease to severe pneumonia and acute respiratory distress syndrome in the human population^1–4^. Many severe COVID-19 patients experience a hyper-inflammatory response (“cytokine storm”)^5^ combined with dysregulated coagulation^6–12^. Both SARS-CoV-2 and the related SARS-CoV-1 coronavirus fuse with and enter cells following engagement of their surface spike (S) protein with human angiotensin-converting enzyme 2 (hACE2)^13–15^, which associates with type II transmembrane serine protease (TMPRSS2) on multiple epi/endothelial tissues and vasculature, including in the lung, liver, colon, esophagus, small intestine, duodenum, kidney, brain, and tongue^16, 17^. TMPRSS2 increases viral uptake by cleaving the S protein, thereby facilitating fusion of the S protein receptor binding domain (RBD) with hACE2 on epi/endothelial cells^13^.

Appropriate, clinically relevant animal models are required to evaluate mechanisms of human-like pathogenesis following SARS-CoV-2 infection and as platforms for rapid testing of vaccines and potential therapeutics. The SARS-CoV-2 S protein has lower affinity for murine (m)ACE2 than for hACE2^15^, and accordingly, mice are not susceptible to infection by coronaviruses that utilize this receptor. To overcome this limitation, several murine strains transgenic (Tg) for hACE2 expression driven by various promoters have been generated to facilitate coronavirus research. Overall, these murine models have shown different tropisms, viral loads and pathologies following infection^18–21^. Hence, a human immune system (HIS)-humanized animal model expressing hACE2 that is permissive for SARS-CoV-2 infection would be a highly appealing model to study mechanisms of viral entry and human-like anti-viral immune responses. Challenges in developing HIS-mouse models that mimic the human immune system with high fidelity include poor engraftment of hematopoietic stem cells, inefficient human cell expansion and homeostasis, insufficient numbers of reconstituted human T or B cells, sub-optimal B-cell development, lack of immunoglobulin class switching, acute/chronic GVHD, and lack of HLA class I and/or II T-cell restriction^22, 23^. Furthermore, restrictions on the use of human fetal tissue for research (https://oir.nih.gov/sourcebook/ethical-conduct/special-research-considerations/policies-procedures-use-human-fetal-tissue-hft-research-purposes-intramural/policies) has focused attention on alternative donor cell sources, such as umbilical cord blood.

The HIS-humanized DRAGA (HLA-DR4.HLA-A2.1.IL-2Rγc KO.RAG1 KO.NOD) mouse strain is devoid of many of the above limitations^24, 25^. These mice are HIS-reconstituted after irradiation by infusion with hematopoietic stem cells from HLA-matched umbilical cord blood. They lack the murine adaptive immune system while expressing a long-lived functional HIS. They respond vigorously by specific T and B cell responses to infection or immunization with various pathogens including malaria protozoans, HIV, ZIKA, Scrub Typhus, and Influenza type A heterosubtypes^26–31^. They also engraft various hematopoietic cell-derived human cells, including endothelial cells (EDs) in the liver^26^ and both epithelial cells (ECs) and EDs in the lungs^28, 29^.

Herein, we demonstrate that HIS-DRAGA mice naturally express hACE2 and hTMPRSS2 on engrafted human epi/endothelial cells on several organs, including the lungs. Importantly, these mice sustained intranasal infection with SARS-CoV-2 for at least 25 days and developed dose-dependent, mild to severe human-like lung immunopathology. We suggest that the HIS-DRAGA mouse is an excellent and convenient model for studying mechanisms of SARS-CoV-2 infection and human-like anti-SARS-CoV-2 immune responses, and for testing the safety and efficacy of novel vaccines and therapeutics.

## Results

### HIS-DRAGA mice naturally express hACE2 and hTMPRSS2

Human *ACE2* mRNA was detected in the lungs of non-infected, HIS-reconstituted DRAGA mice (mice ***a-j***, **Table S1**) but not of non-reconstituted mice, whereas *mACE2* mRNA was detected in both HIS-reconstituted and non-reconstituted DRAGA mouse lungs (**Fig. 1a**). A recombinant SARS-CoV-2 S1(RBD)-mouse Fcγ2a chimeric protein (S1(RBD)-mFcγ2a) and magnetic beads coated with rat anti-mouse IgG2a were used to immunoprecipitate hACE2 from pooled lung homogenates of HIS-reconstituted DRAGA mice (n=10), non-HIS reconstituted DRAGA mice (n=10) and a human lung tissue lysate. Quantitative hACE2 ELISA measurements indicated hACE2 was 7.8X less abundant in HIS-DRAGA mouse lungs than in the human lung sample, while no hACE2 was detected in immunoprecipitates from non-HIS reconstituted DRAGA mouse lungs (**Fig. S1**). The molecular weight of hACE2 expressed in lungs of HIS-DRAGA mice was identical to that in human lungs as shown by Western Blot (**Fig. S1**). To visualize hACE2 protein expression, immunofluorescence microscopy was carried out on lung sections probed with S1(RBD)-mFcγ2a. Digitized images revealed hACE2 expression on alveolar and bronchiolar epithelia (**Fig. 1b**), while very weak or no staining was detected in lung sections from non-HIS reconstituted mice (**Fig. 1c)**. Immunofluorescence staining also revealed that hTMPRSS2 was co-localized with hACE2 on the bronchiolar epithelia and on endothelial walls of pulmonary arterioles (**Fig. 1d**).

**Figure 1.**
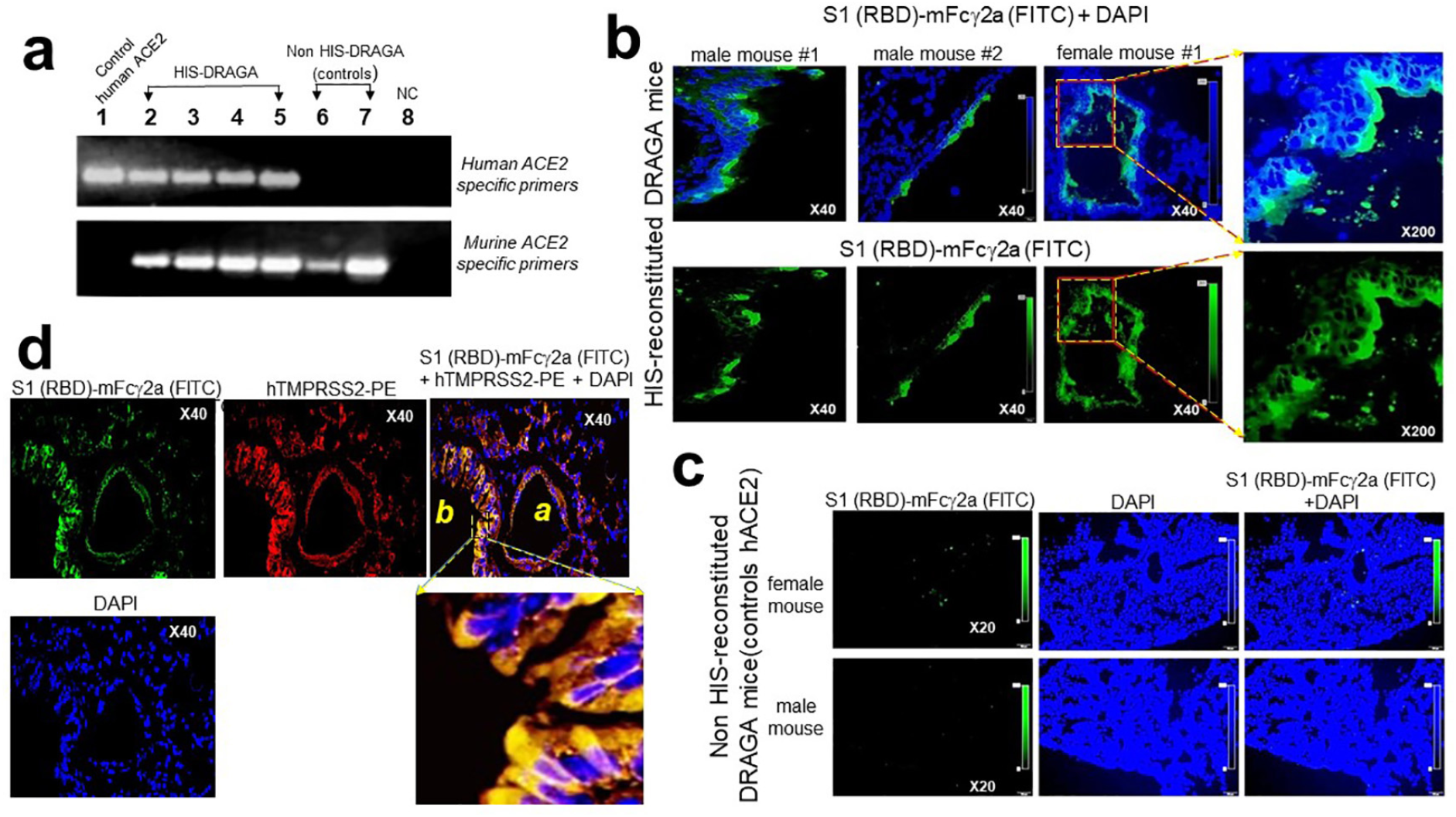
Human ACE2 and TMPRSS2 detection in the lungs of non-infected HIS-DRAGA and DRAGA mice. **a.** Positive control (+) = human lung mRNA. Negative control (−) = primers alone. *upper row*, PCR amplicons of *hACE2* from the lungs of 4 HIS-DRAGA mice (2 females and 2 males) and 2 non-HIS-reconstituted DRAGA mice (1 female and 1 male), amplified using hACE2-specific primers. *Lower row,* PCR amplicons of the same lung samples in the upper row, amplified using mACE2-specific primers. **b.** hACE2 protein expression on alveolar epithelia as indicated by binding of the S1(RBD)-mFc□2a protein + goat anti-mouse IgG-FITC in lung sections from a representative HIS-DRAGA female mouse and two HIS-DRAGA male mice. *Upper panels,* merged images of S1(RBD)-mFc□2a binding (green) and nuclei (blue, DAPI), and an enlargement of the hACE2^+^ alveolar epithelia. *Lower panels*, binding of S1(RBD)-mFc□2a protein + goat anti-mouse IgG-FITC and an enlargement of the hACE2^+^ alveolar epithelia. **c.** Same staining protocol as in **a** for lung sections from two non-HIS reconstituted DRAGA mice showing very weak binding (female mouse) and no detectable binding (male mouse) of S1(RBD)-mFc◻2a protein, indicating siginificantly weaker avidity of the S1 viral protein for the murine than the human ACE2 receptor. **d.** lung section from a representative HIS-DRAGA female mouse showing hTMPRSS2 co-localization with hACE2 as revealed by S1(RBD)-mFc◻2a protein + goat anti-mouse IgG-FITC (green) and anti-human TMPRSS2-PE (red) on the epithelial wall of a bronchiole (*b*) and the endothelial wall of a pulmonary arteriole (*a*). A section of the bronchiole image is also shown enlarged 200X.

We next questioned whether human (h)CD326, a specific marker of lung ECs, is also expressed on engrafted hECs. Co-staining with anti-hCD326-PE and S1(RBD)-mFcγ2a revealed co-localization of hACE2 and hCD326 in lungs of HIS-reconstituted (**Fig. S2,** *upper* and *middle panels*) but not non-reconstituted DRAGA mice (**Fig. S2,** *lower panels*). Staining of sections from liver, kidneys, small intestine and brain of HIS-DRAGA mice with S1(RBD)-mFcγ2a revealed hACE2 expression on liver endothelia (**Fig. S3**), epi/endothelia of proximal and distal convoluted tubules and small arterioles surrounding the podocytes in the kidney glomeruli (**Fig. S4**), columnar epithelia of the absorptive intestinal villi (**Fig. S5**), and the white matter, granular layer and Purkinje cells in the cerebellum cortex (**Fig. S6**). Expression of hACE2 on epi/endothelia of these organs was further confirmed by probing the tissue sections with a mouse anti-hACE2 antibody (**Fig. S7**).

### HIS-DRAGA mice can sustain SARS-CoV-2 infection

In the first infection experiment, three HIS-DRAGA mice were infected intranasally with relatively high doses of virus in 50 μl saline, 25 μl per nostril: 2.8×10^4^ pfu (male #M1 + female #F1) and 2.8×10^3^ pfu (female #F2). While the male mouse succumbed 24 hours after infection, both female mice sustained the infection until the experimental endpoint of 14 days post-infection (dpi) (**Fig. 2a**). These mice showed an abrupt loss in body weight (likely due to severe dehydration), ruffed fur, hunched back, and reduced mobility starting at 1 dpi. Mouse #F2 regained its original weight and mobility by 9 dpi, while mouse #F1 was still 10% below its original weight at 14 dpi, when they were both euthanized.

**Figure 2.**
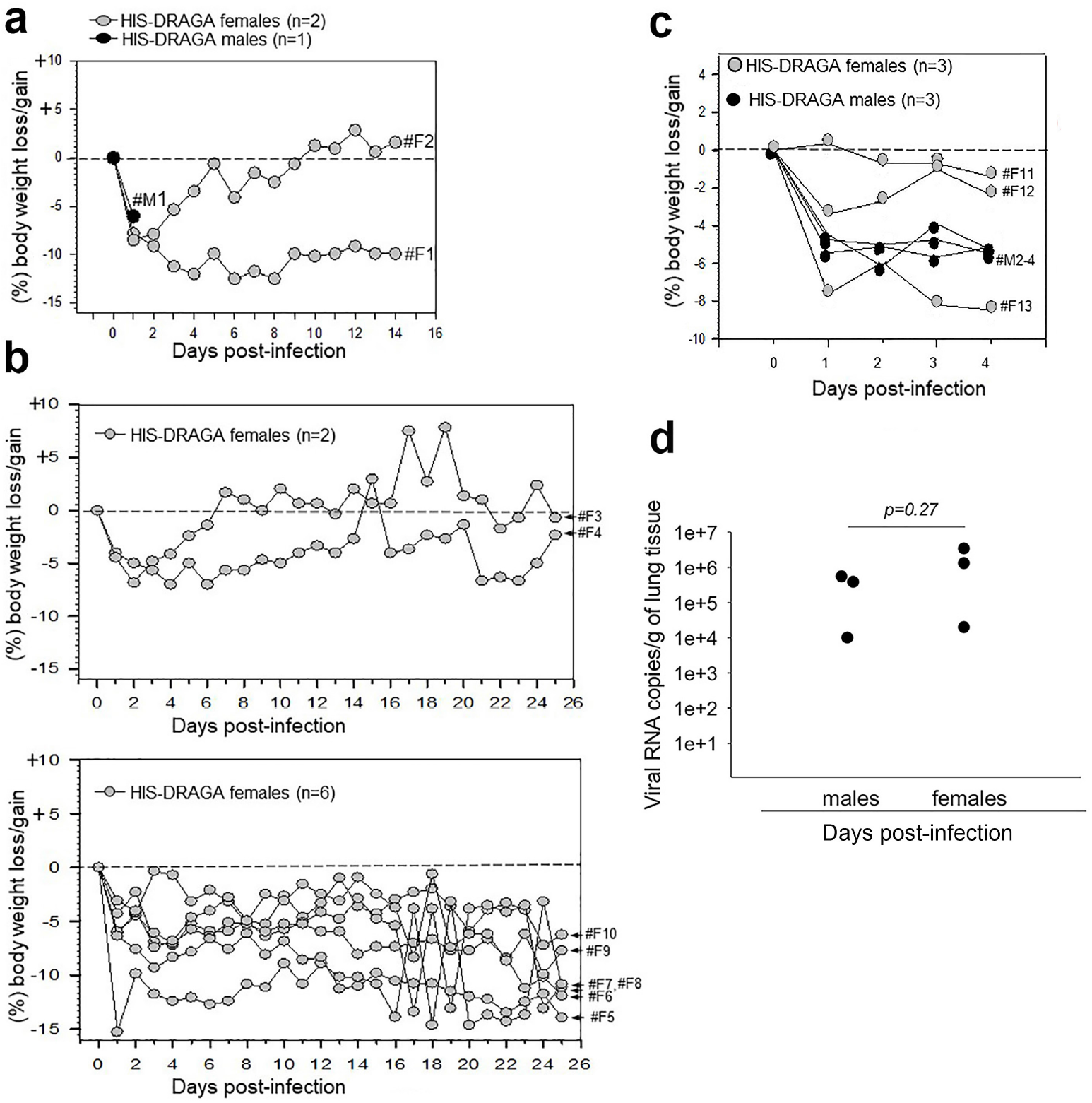
Post-infection body weights and viral titers. **a.** Changes in % body weights of HIS-DRAGA mice #M1 infected with SARS-CoV-2 virions (2.8×10^4^ pfu) and #F1 and #F2 infected with 2.8×10^4^ pfu and 2.8×10^3^ pfu, respectively (1^st^ infection experiment). **b.** Changes in % body weights for 8 HIS-DRAGA mice infected with 10^3^ pfu of SARS CoV-2 virions (2^nd^ infection experiment). Shown are the mice that recovered their initial body weights after infection (upper panel), and those that did not (lower panel). **c.** Changes in % body weights for 6 HIS-DRAGA mice infected with 10^3^ pfu SARS-CoV-2 virions (3^rd^ infection experiment). **d.** SARS-CoV-2 viral loads in the lungs of 6 infected HIS-DRAGA mice (10^3^ pfu/mouse) as quantified by RT-qPCR (3^rd^ infection experiment) at 4 dpi. There was no significant difference between the viral RNA copies in lungs of female versus male mice.

In the second infection experiment, 8 HIS-DRAGA female mice (#F3-F10) were inoculated i.n. with a lower dose of SARS-CoV-2 virus (10^3^ pfu/mouse), and their body weights and clinical condition were monitored daily for 25 days (**Fig. 2b**). All mice in this group lost 5%-15% of their weight and showed deteriorating conditions (ruffed fur, hunched back, reduced mobility) starting at 1 dpi. However, mice #F3 and #F4 recovered their initial body weights by 7 and 25 dpi, respectively, while the remaining mice survived the infection but had still not recovered their initial body weights at the experimental endpoint (25 dpi), when they were euthanized for further analyses.

In the third infection experiment, 3 HIS-DRAGA females and 3 HIS-DRAGA males (#F11-F13 and #M2-M4, **Table S1**) were infected i.n. with SARS-CoV-2 (10^3^ pfu/mouse) and their body mass and clinical condition were monitored daily. Five of these 6 mice showed weight loss starting at 1 dpi (**Fig. 2c**) and all showed ruffed fur, hunched back and reduced mobility. The mice were sacrificed at 4 dpi, and the viral loads in their lungs were quantified (**Fig. 2d**).

### SARS-CoV-2 infected HIS-DRAGA mice display severe, human-like lung pathology

Patients infected with SARS-CoV-2 have been shown to exhibit varying degrees of lung pathology that correlate with the viral load in their lungs^1–4^. As in recent reports based on human autopsies^32–34^, and in contrast to the non-infected mice (**Fig. 3a**), H&E and Masson’s trichrome staining of lung sections from mice infected with high doses of SARS-CoV-2 virions showed multiple interstitial, peripheral, and peri-bronchiolar infiltrates (**Fig. 3b**). These were more noticeable in the lungs of mouse #F1, which was infected with the highest dose of virus (2.8×10^4^ pfu) and had not recovered its initial body weight by 14 dpi than in mouse #F2, which was infected with 2.8×10^3^ pfu, (**Fig. S8**). Mouse #F1 also developed multiple intra-alveolar and intra-arteriolar microthrombi adherent to the endovasculature (**Fig. 3c, d**) that stained positive for the platelet marker CD61 (glycoprotein IIIa) (**Figs. S9** and **S10**), as well as intra-bronchiolar blood clots (**Fig. 3e**) and interstitial hemorrhagic patches (**Fig. 3f**).

**Figure 3.**
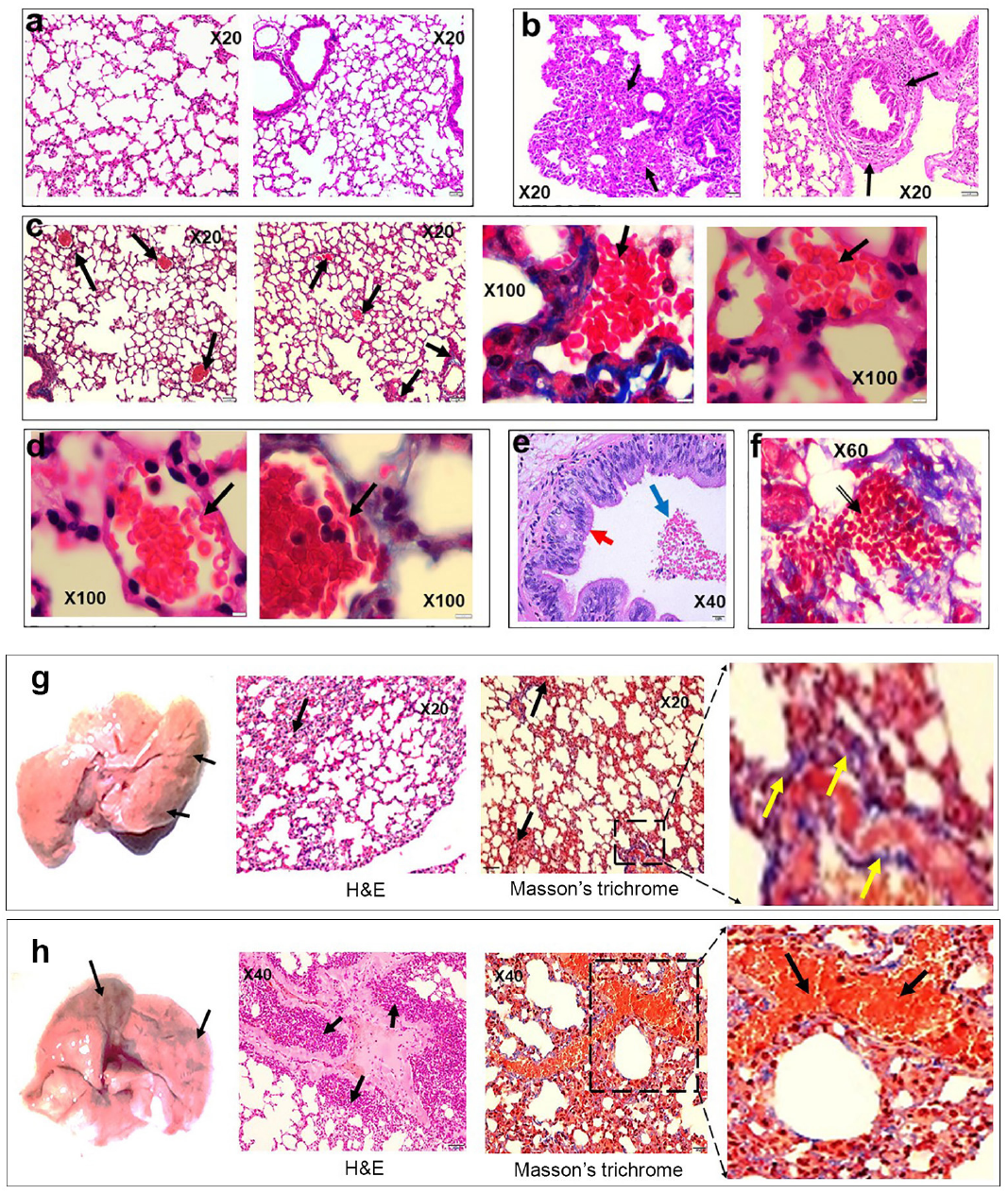
Lung pathology of HIS-DRAGA mice infected with SARS-CoV-2. **a.** H&E-stained lung sections from two non-infected HIS-DRAGA mice showing healthy alveolar architecture and two bronchioles lacking lymphocytic infiltration. **b.** Two separate H&E-stained sections of the lungs from HIS-DRAGA mouse #F1 infected with 2.8×10^4^ pfu, which did not recover its initial body weight at the experimental endpoint (14 dpi). Heavily infiltrated areas at the lung periphery and surrounding a bronchiole are shown (arrows), and distorted alveolar architecture was also observed. **c.** Masson’s Trichrome-stained sections of a lung from mouse #F1 showing multiple intra-arteriolar microthrombi (left panel, arrows) and intra-alveolar microthrombi (3 right panels, arrows). **d.** Masson’s Trichrome-stained sections of a lung from mouse #F1 showing adherence of red blood cells to the endovasculature (arrows). **e.** H&E staining of a representative lung section from mouse #F1 showing a large intra-bronchiolar blood clot (blue arrow). Also shown are the bronchus columnar ciliated cells of epithelia (red arrow). **f.** Representative Masson’s Trichrome-stained section of a lung from mouse #F1 showing a large interstitial hemorrhagic area (arrow). **g.** Morphologic image of the lungs (left panel) and H&E and Masson’s Trichrome-stained sections of the lungs (right panels) from the HIS-DRAGA mouse #F3 infected with 10^3^ pfu, which recovered its initial body weight at 7 dpi. Discoloration on the left lobe (arrows) and a few, small interstitial and peripheral infiltrates are indicated by arrows. Enlargement shows incipient peri-alveolar collagen deposition as a sign of developing pulmonary sequelae. **h.** Morphologic image of the lungs (left panel) and the H&E and Masson’s Trichrome-stained sections of the lungs (*right panels*) from HIS-DRAGA mouse #F5 infected with 10^3^ pfu of SARS-CoV-2, which sustained the infection but did not recover its initial body weight by 25 dpi. Shown is a large discolored area of the upper right lobe (arrow), heavy infiltration with disrupted alveolar architecture (arrows, *middle panel*), and a large hemorrhagic patch in vicinity of a major bronchiole (arrows, *right panel* and *enlargement*).

Lung tissues from female HIS-DRAGA mice infected i.n. with the lowest dose of virus (10^3^ pfu/mouse, second experiment) were analyzed histologically at the experimental endpoint (25 dpi). Morphological analysis of lungs from mice #F3 and #F4, which recovered their initial body weights by 7 and 25 dpi, respectively, showed small discolored areas and dispersed interstitial infiltrates throughout the lung parenchyma (**Fig. 3g**). In contrast, the mice that had not recovered their initial body weights by 25 dpi, e.g., mouse #F5, showed large pulmonary discoloration and heavily infiltrated areas (**Fig. 3h**) and incipient collagen-based fibrosis in peri-alveolar infiltrated areas (**Figs. 3g** and **S10**). Similar to mouse #F1 that was infected with 2.8×10^4^ pfu in the first experiment (**Fig. 3f**), mouse #F5 showed interstitial hemorrhagic patches in the lungs (**Fig. 3h**) and a more advanced process of pulmonary fibrosis (**Fig. S11**).

### Pulmonary infiltrates in SARS-CoV-2 infected HIS-DRAGA mice contained lung-resident human T cells expressing activation markers

Infiltrates visualized in lung sections from SARS-CoV-2-infected mice stained positive for hCD45 indicating human lymphocytes (**Fig. 4a,c**) and hCD3 indicating human T cells (**Fig. 4b,d**), as we previously described in non-infected HIS-humanized DRAGA mice^24, 25^. Interestingly, some of the human CD3^+^ T-cell infiltrates in mice infected with a relatively high SARS-CoV-2 dose (2.8×10^3^ pfu) looked fairly organized (**Fig. 4d**). Both hCD8^+^ and hCD4^+^ T-cell subsets were found sequestered in alveolar CD326^+^ hECs, with some egressing into the alveolar air space (**Fig. 5**). Alveolar hCD326^+^ hECs contained larger clusters of hCD8^+^ T cells than hCD4^+^ T cells. Human CD8^+^ T cells that co-localized with hCD326^+^ ECs also stained positive for CD103, a marker for lung T-cell residency^35, 36^ (**Fig. 6).** Among the CD8^+^ T cells sequestered in alveolar EC niches, many stained positive for perforin and granzyme B, indicating potential cytotoxic activity **(Fig. 6**).

**Figure 4.**
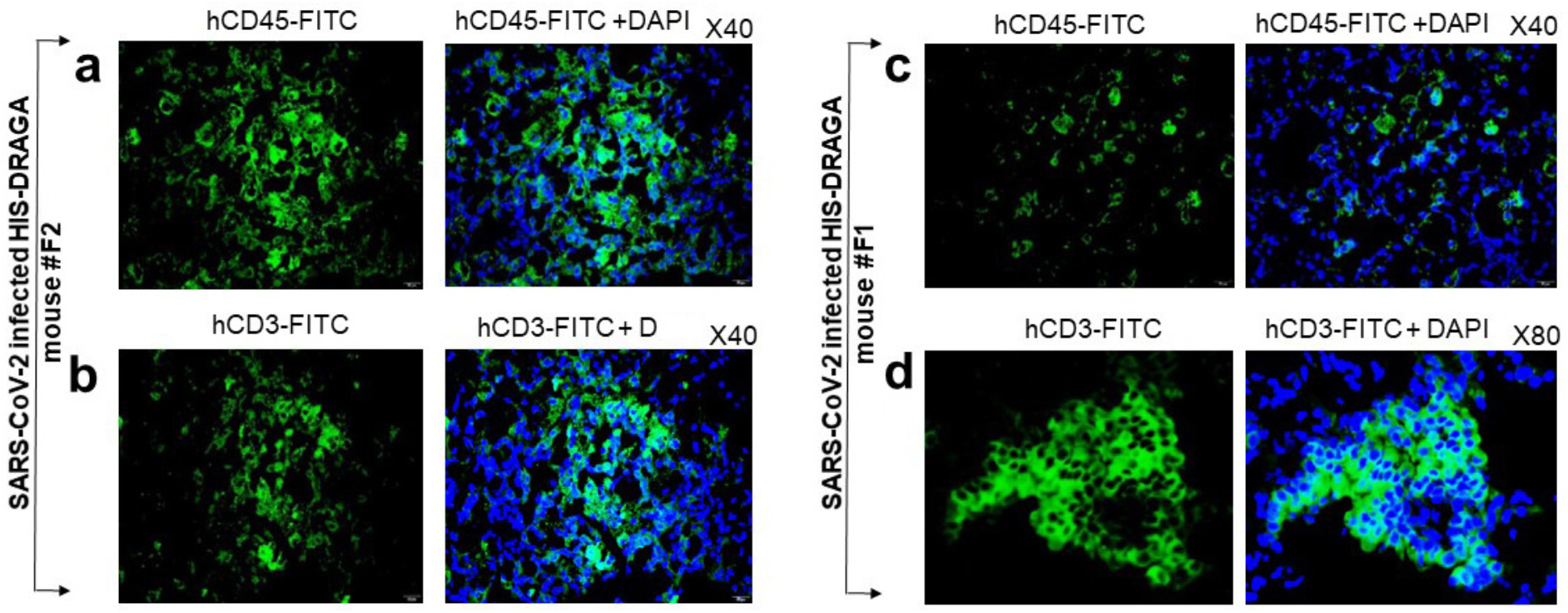
Infiltrating human lymphocytes in the lungs of SARS-CoV-2 infected HIS-DRAGA mice. Lung sections from HIS-DRAGA mice #F2 and #F1 infected with 2.8×10^3^ pfu and 2.8×10^4^ pfu, respectively, stained with anti-hCD45-FITC or anti-hCD3-FITC at the experimental endpoint (14 dpi). **a,b**. hCD45^+^ and hCD3^+^ cell infiltrates in the lungs of #F2 mouse recovering from infection. **c,d.** hCD45^+^ and hCD3^+^ cell infiltrates in the lungs of mouse #F1, which did not recover from infection. Of note, infiltrating CD3^+^ T cells were clustered in lung epithelial niches of mouse #F2 whereas CD3^+^ T cell infiltrates in mouse #F1 were more dispersed.

**Figure 5.**
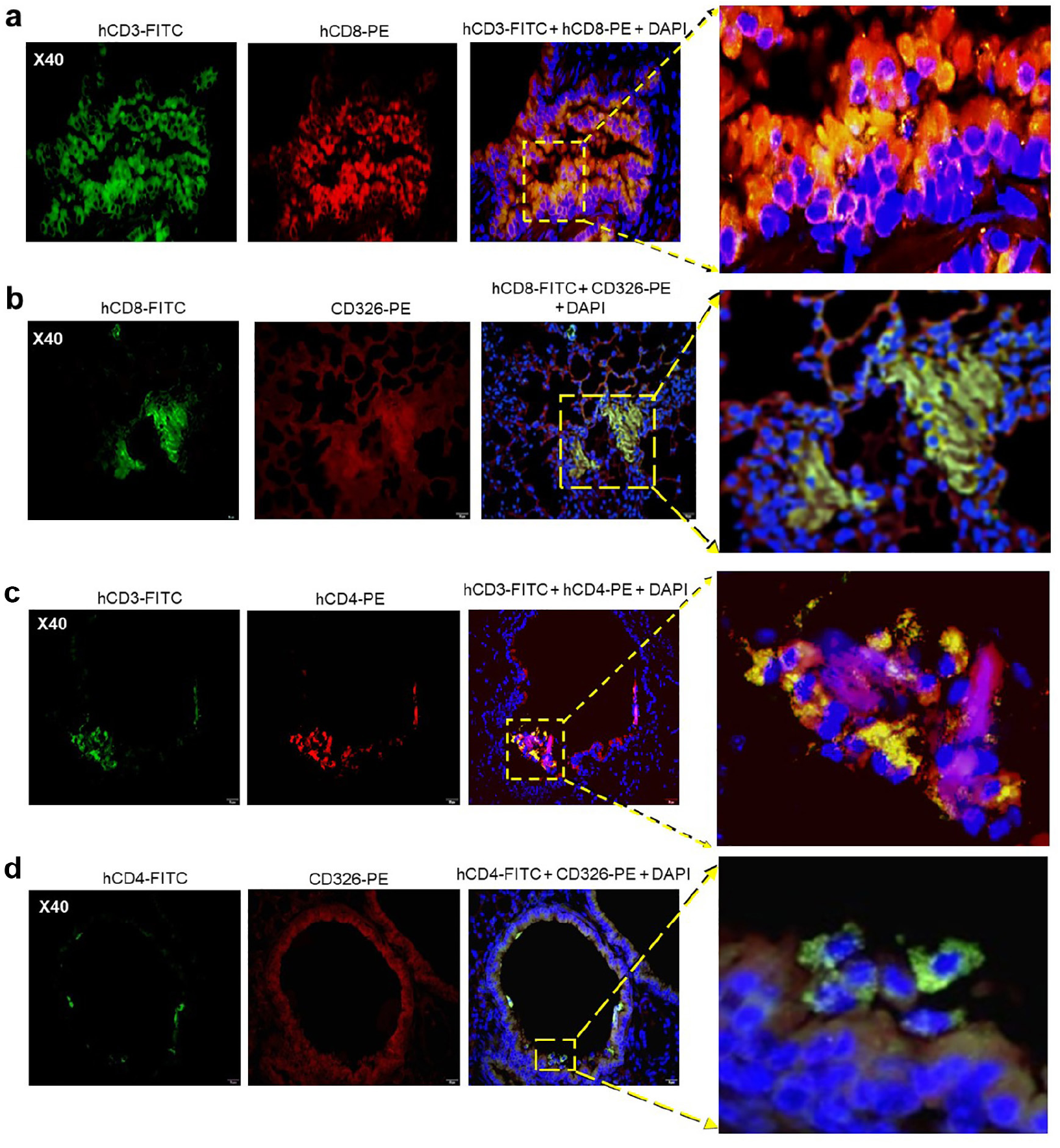
Human CD8^+^ and CD4^+^ T cells in CD326^+^ lung niches in SARS-CoV-2 infected HIS-DRAGA mice. **a.** Representative lung section from HIS-DRAGA mouse #F2 infected with 2.8×10^3^ pfu that recovered its initial body weight at 14 dpi, co-stained with anti-hCD3-FITC and anti-hCD8-PE antibodies. Enlargement shows an area with a large cluster of hCD3^+^CD8^+^ T cells (orange) sequestered in the lung epithelia, with some hCD3^+^CD8^+^ T cells egressing into the alveolar air space. **b.** Lung section from the same infected mouse (#F2) co-stained for hCD8 and hCD326 (epithelial marker). Enlargement shows a large cluster of hCD8^+^ T cells co-localized with the hCD326^+^ lung epithelia. **c.** Lung section from HIS-DRAGA mouse #F2 infected with 2.8×10^3^ pfu that recovered its initial body weight at 9 dpi, co-stained with anti-hCD3-FITC and anti-hCD4-PE antibodies. Enlargement shows a few hCD3^+^CD4^+^ T cells (yellow). **d.** lung section from the same mouse (#F2) showing hCD4^+^ T cells co-localized with the CD326^+^ epithelial layer. Enlargement shows an area containing co-localized CD326^+^ lung ECs with CD4^+^ T cells egressing into the alveolar air space (orange).

**Figure 6.**
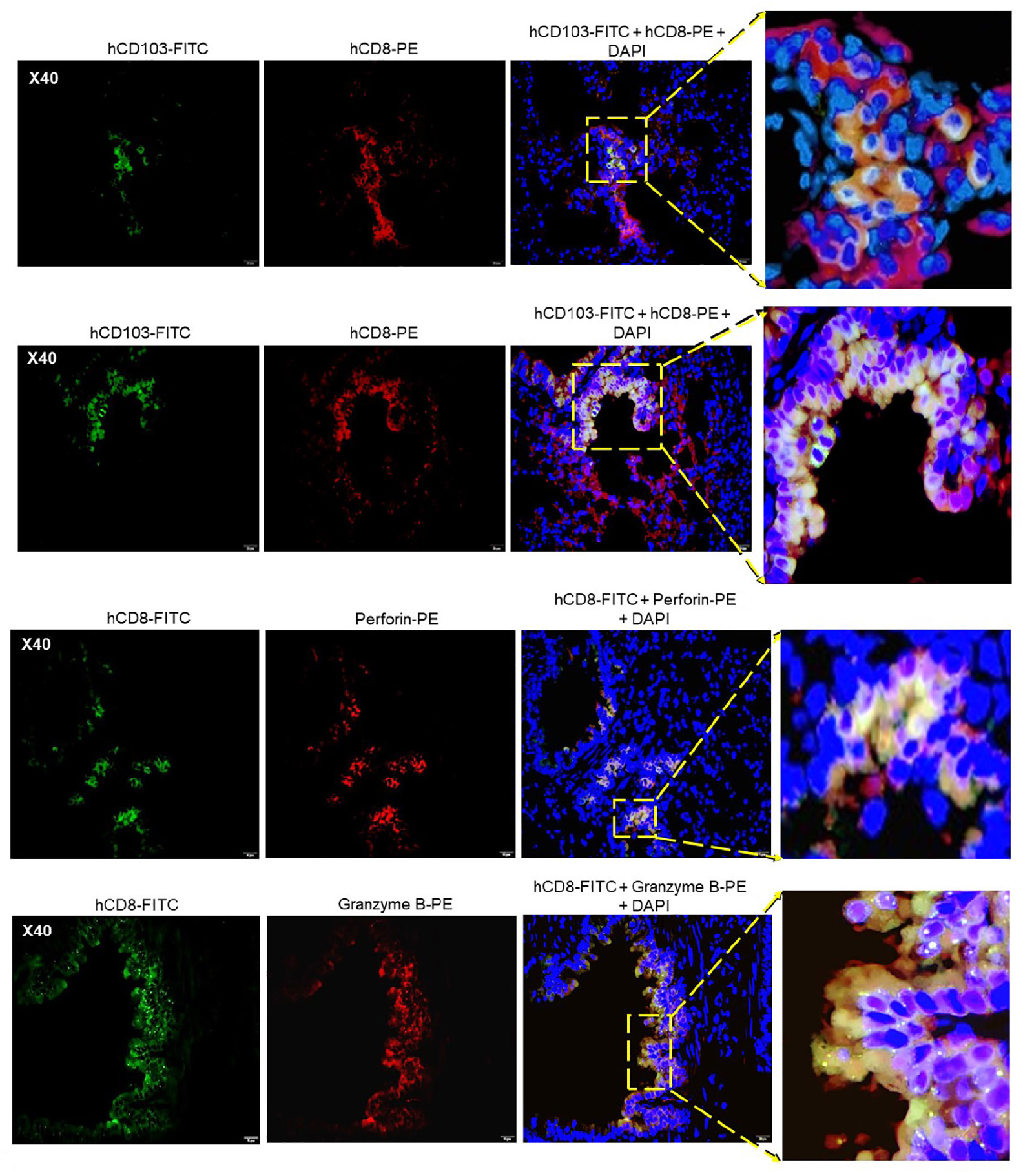
hCD8^+^ T-cell residency and cytotoxicity in the lung epithelial niches of a SARS-CoV-2-infected HIS-DRAGA mouse. Shown are representative lung sections from mouse #F2 infected with 2.8×10^3^ pfu, which recovered its initial body weight at 9 dpi. **a**. Lung section co-stained with anti-hCD8-PE and anti-hCD103-FITC (marker of lung residency). Enlargement shows a cluster of hCD8^+^ hCD103^+^ T cells sequestered in a lung alveolar niche. **b**. Lung section co-stained with the same antibodies. Enlargement shows several large epithelial niches containing hCD8^+^hCD103^+^ T cells (orange), with some cells (orange) egressing into the alveolar air space. **c**. hCD8^+^ T cells clustered in the alveolar epithelial niches staining positive for perforin (orange). **d.** hCD8^+^ T cells from the same mouse (#F2) sequestered throughout the alveolar epithelial niches and staining positive for Granzyme B (orange), with some egressing into the alveolar air space.

### HIS-DRAGA mice infected with SARS-CoV-2 mounted specific antibody responses to SARS-CoV-2 viral proteins

ELISA results obtained using plates coated with recombinant SARS-CoV-2 S1(RBD), S1-trimer or N proteins were used to measure specific antibodies in serum samples derived from infected mice. Mouse #F2 from the first experiment, which was infected with 2.8×10^3^ pfu and recovered its initial body weight by 9 dpi, developed lung-resident (CD103^+^) hCD8^+^ T cells (**Fig. 6**) but did not have detectable S1(RBD)-specific hIgM antibodies at 14 dpi (not shown). In contrast, mouse #F1, which was infected with 2.8×10^4^ pfu and had not recovered its body weight by 14 dpi, developed S1(RBD)-specific hIgM antibodies, but had fewer clusters of lung-resident hCD8^+^ T cells (not shown). Furthermore, the second infection experiment using a lower dose of SARS-CoV-2 virions (10^3^ pfu) and a longer follow-up period (25 dpi), revealed that all mice developed hIgM and hIgG antibody responses to the spike and nucleocapsid (N) viral proteins (**Fig. S12**). Of note, the mice that recovered their initial body weights, #F3 and #F4, mounted the strongest hIgG responses to S1(RBD), S-trimer and N viral proteins.

Together, these results have demonstrated that pluripotent human CD34^+^ human stem cells from umbilical cord blood differentiated not only into functional human T and B cells, but also into human ECs and EDs expressing hACE2 and hTMPRSS2 in several organs of HIS-DRAGA mice. HIS-DRAGA mice sustained infection with SARS-CoV-2 for at least 3 weeks, exhibiting clinical symptoms, with some recovering from the infection. Evaluations of their organ tissues have also revealed pathological events including parenchymal and peripheral lung CD3^+^ T-cell infiltrates, intra-alveolar and intra-arteriolar microthrombi adherent to the endovasculature, intra-bronchiolar blood clots and incipient pulmonary sequelae. Lung-resident (CD103^+^) CD8^+^ secreting granzyme B and perforin, and CD4^+^ T cells have been visualized in CD326^+^ epithelial lung niches of infected HIS-DRAGA mice, as previously shown in influenza-infected human lung tissue^35, 36^.

## Discussion

The HIS-humanized DRAGA mouse expresses a sustained, fully functional human immune system. In the present study, we found that reconstituted HIS-DRAGA mice engraft human epi/endothelial cells expressing hACE2, the primary receptor for SARS-CoV-2 and its associated serine protease, hTMPRSS2, in multiple organs. Recent studies have revealed that ACE2 is widely expressed in human tissues^37^.

We have demonstrated that HIS-DRAGA mice can be readily infected via the intranasal route with non-mouse-adapted SARS-CoV-2 and that the infection is sustained for at least 25 days. Infected mice developed mild to severe lung immunopathology including T-cell infiltrates, alveolar damage, intra-alveolar and intra-arteriolar microthrombi, bronchiolar blood clots and collagen deposition in the lungs. Severely infected HIS-DRAGA mice recapitulated many of the diverse pathological events in COVID-19 patients involving lungs^1–4, 38^, liver^39^, gastro-intestinal track^40^, kidneys^41^, central nervous system^42^, heart^43^ and genitourinary system^44^. As in recent analyses of autopsy samples from COVID-19 patients^32^, the lung infiltrates of infected HIS-DRAGA mice were interstitial and grouped around terminal bronchioles and peripheral parenchyma. They contained lung-resident (CD103^+^) human T cells, particularly CD8^+^ T cells clustered in CD326^+^ lung epithelial niches that stained positive for granzyme B and perforin, indicating the presence of functional cytotoxic T cells. The presence of CD4^+^ and CD8^+^ T cells reactive to the ORF-1 and NP proteins of SARS-CoV-2 virus in convalescent COVID-19 patients was reported recently^45^. Some granzyme^+^ and perforin^+^ lung-resident CD8^+^ human T cells visualized in the infected HIS-DRAGA mice egressed into the alveolar air space. Lung-resident CD8^+^ T cells have been detected and characterized in mice, monkeys and humans^46, 47^. This cell subset is positioned to act at the frontline of lung epithelial mucosa after a primary viral respiratory infection to provide rapid and efficient cross-protection against subsequent exposures to respiratory viruses^46–51^.

Interestingly, clusters of lung-resident CD8^+^ T cells were more profuse in the infected mouse (#F2) that recovered its body weight early after infection with a relatively high dose of CoV-2 virions (2.8×10^3^ pfu) but showed no detectable S1(RBD)-specific hIgM at 14 dpi. In contrast, mouse #F1, which was infected with 2.8×10^4^ pfu and had not recovered its body weight at 14 dpi, developed S1(RBD)-specific hIgM antibodies and showed fewer human lung-resident T cells. Furthermore, two of the mice infected with 10^3^ pfu recovered their initial body weights by 7 and 25 dpi (#F3 and #F4, respectively), and these mice developed the highest titers of hIgG antibodies to the Spike(S) and Nucleocapsid (N) proteins. They also showed more profuse clusters of lung-resident (CD103^+^) hCD8^+^ T cells (not shown) as compared with mice that did not recover from the infection. These findings raised a question that needs to be further addressed: are antibodies, lung-resident CD8^+^ T cells, or both critical for efficient protection against COVID-19? Recent studies in animal models and humans have suggested critical roles for both antibodies^52–54^ and T cells^55^ for protection against COVID-19. Another subject for future research is the possibility of natural transmission of SARS-CoV-2 infection among mice. We recently demonstrated such transmission of influenza between co-caged, infected and non-infected HIS-DRAGA mice^31^. While investigation of the dynamics of antibody titers and virus clearance in COVID-19 patients is feasible in humans, exploring the dynamics of lung-resident CD8^+^ T cells in infected human subjects is not, as studies of human lungs are restricted to analyses of single biopsy/resection or post-autopsy samples.

We also found that the SARC-CoV-2 infected HIS-DRAGA mouse are able to mount fully human IgM and IgG antibody responses to the viral proteins, e.g., S1(RBD), trimeric S and N proteins, some of which may have translational potential. While high antibody titers to the S1(RBD) viral protein correlated with body weight recovery following infection, i.e. with less severe infection, the antibody titers to the Nucleocapsid (N) protein did not.

Thrombophilia, including microthrombi in the lung, sometimes termed “immunothrombi” due to their association with the hyperinflammatory response, are a feature of severely infected COVID-19 patients. Histologic analysis of pulmonary vessels from COVID-19 autopsy samples showed widespread thrombosis with microangiopathy^6, 7, 9, 10^. In lungs, new vessel growth occurs predominantly through a mechanism of intussusceptive angiogenesis^56^. In addition, occlusion of alveolar capillaries described in patients with severe COVID-19^11^ and adherence of microthrombi to the vascular endothelium suggest a distinctive angiocentric feature^6, 11^. Lungs of infected HIS-DRAGA mice had clusters of non-nucleated cells staining positive for CD61, consistent with the presence of platelets and microthrombi in the alveolar air space. Infected mice also showed hemorrhagic patches in the lungs, mimicking those described in autopsy samples from humans exiting COVID-19^32^.

Pulmonary fibrosis, also known as sequelae, indicates tissue scarring during healing that occurs by deposition of collagen in heavily infiltrated areas, as revealed in lungs from patients recovering from severe respiratory infections. Radiographic and autopsy data have identified pulmonary fibrosis not only in COVID-19, but also in SARS CoV-1 and MERS^57^. It has been suggested that collagen deposition in severe lung injury by SARS viruses relies primarily on trafficking circulating fibrocytes to the lung^58^ and increased signaling through the epidermal growth factor receptor^59^. Masson’s Trichrome staining of lung sections from SARS-CoV-2-infected mice that survived high-viral-dose infection, and from some mice that sustained low-dose infection for up to 25 days, revealed incipient collagen depositions in heavily infiltrated alveoli and surrounding the bronchioles.

In summary, HIS-DRAGA mice, infected with a non-mouse adapted strain of SARS-CoV-2 were able to sustain the infection for at least 25 days, recapitulated major immunopathological events described in COVID-19 patients, and mounted human cellular and humoral responses to the SARS-CoV-2 proteins. This HIS-humanized *in vivo* surrogate mouse model offers compelling advantages for studying the mechanisms of SARS-CoV-2 infection and human immunopathology of COVID-19 disease, notably the ability to analyze physiological responses and harvest tissues at specific time points following infections and subsequent viral challenges. This mouse model should also prove useful for efficient preclinical testing of both safety and efficacy of vaccines and potential therapeutics for human COVID-19.

## Data and animal availability

Additional datasets generated and/or analyzed during the current study are available from the corresponding authors on reasonable request.

## Acknowledgements

We acknowledge support from the Collaborative Health Initiative Research Program at Uniformed Services University, Bethesda, MD (K.P.P), the Department of Medicine, Sanford Endowment intramural funds (T-D.B. and K.P.P.), the Military Infectious Diseases Research Program (S.C.), and NHLBI HL 130448 and HL 12767B (K.P.P).

We thank Soumya Sashikumar for maintaining DRAGA mice colony at NMRC. The study protocol was reviewed and approved by the Bioqual Institutional Animal Care and Use Committee (#20-019P) and by the Walter Reed Army Institute of Research/Naval Medical Research Center Institutional Animal Care and Use Committee (#19-IDD-24) in compliance with all applicable federal regulations governing the protection of animals and research. T-D.B, KPP, KKC and SC are federal employees and this work was conducted as part of their official duties.

*The opinions or assertions contained herein are the private ones of the authors and are not to be construed as official or reflecting the views of the Department of Defense or the Uniformed Services University of the Health Sciences.*

## Author contributions

T-DB, SC, KPP and KKC conceived and guided the study. T-DB, SC, PV and AFK performed experiments. T-DB, SC and KPP analyzed data and wrote the paper, with input from PV and AFK. SK, DB, ML, JG, TP-T and CK oversaw or conducted the experiments at the BSL-3 facility at Bioqual, Inc. (Rockville, MD, USA): infecting and monitoring the clinical condition of mice, extracting lung tissues, and measuring lung viral loads by RT-qPCR. All authors approved the content of the manuscript.

## ONLINE METHODS

### HIS-reconstitution of DRAGA mice

DRAGA mice express the HLA-A2.1 and HLA-DR0401 transgenes on a Rag1KO.IL2RγcKO.NOD (NRG) background, and they have been described previously^28–31^ HLA-A2.1.HLA-DR0401 positive umbilical cord blood was obtained from the New York Blood Center (Long Island City, NY, USA). Mice were irradiated (350 rads) and injected intravenously with CD3^+^ T-cell-depleted cord blood cells (EasySep Human CD3 Positive Selection Kit, Stem Cell Technologies, Vancouver, BC, Canada) containing approximately 10^5^ human CD34^+^ hematopoietic stem cells (HSC) determined by FACS using a mouse anti-human CD34 antibody (BD Biosciences, San Jose, CA, USA) as described^49,52,54^. The procedures for assessing human T and B cell reconstitution in peripheral blood by FACS have been described^49,52,54^. As documented in our previous studies, >90% of HIS-reconstituted DRAGA mice generated using these procedures reconstitute a human immune system by 3 to 4 months post-CD34^+^ HSC infusion. The human reconstitution status of DRAGA mice at the time of our SARS-CoV-2 infection experiments was determined based on FACS measurement of T cells and B cells in peripheral blood (**Table S1**).

### RT-PCR detection of hACE2 mRNA in HIS-DRAGA mouse lungs

RNA was extracted using a Qiagen RNA extraction kit (Qiagen, Hilden, Germany) from lungs of HIS-DRAGA and control (non-HSC-infused DRAGA) mice. Human lung mRNA (Sigma-Aldrich, St. Louis, MO, USA) served as a positive control. PCR primers specific for hACE2 were: forward, CAGGAAATGTTCAGAAAGCA and reverse, TCTTAGCAGAAAAGGTTGTG. The murine ACE2 specific primers were: forward: AGCAGATGGCCGGAAAGTTG, and reverse: TCTTAGCAGGAAAGGTTGCC. RT-PCR was performed using a One-step RT-PCR kit (Qiagen) for 45 cycles using 1 μg RNA template and 1.6 μM of each primer, following the manufacturer’s instructions. The PCR amplicons were run on a 3% agarose gel. PCR bands were purified from the agarose gels and nucleotide sequenced (Eurofins, Coralville, Iowa, USA).

### RT-qPCR measurement of viral RNA copies in SARS-CoV-2 infected HIS-DRAGA mouse lungs

RNA from lungs of mice #M2-M4 and #F11-F13 was extracted using RNA-STAT 60 extraction reagent (Tel-Test, Inc., Friendswood, TX, USA) + chloroform, precipitated and resuspended in RNAse-free water. Control RNA was isolated from SARS-CoV-2 viral stocks following the same procedure and quantified by OD260. These control stocks were serially diluted and OD260 values measured to generate a standard curve. RT-qPCR of the lung RNA was carried out using the following primers : 2019-nCoV_N1-F :5’-GAC CCC AAA ATC AGC GAA AT-3’; 2019-nCoV_N1-R: 5’-TCT GGT TAC TGC CAG TTG AAT CTG-3’; and probe 2019-nCoV_N1-P: 5’-FAM-ACC CCG CAT TAC GTT TGG TGG ACC-BHQ1-3’ (Integrated DNA Technologies, Coralville, IA, USA) which were designed to bind to and amplify a conserved region of SARS-CoV-2 Nucleocapsid (N) RNA. Amplification was performed with an Applied Biosystems 7500 Sequence detector using the following program: 48°C for 30 minutes, 95°C for 10 minutes followed by 40 cycles of 95°C for 15 seconds, and 1 minute at 55°C. Reactions were carried out using a TaqMan RT-PCR kit (Meridian Bioscience, Memphis, TN, USA) in 50 μL volume containing 5 μL of template, 2 μM of each primer and 2μM of each probe. The number of viral RNA copies per mL was calculated by extrapolation from the standard curve, and values were then converted to the number of viral RNA copies per gram of lung tissue.

### Extraction and quantification of hACE2 protein in HIS-DRAGA and non-HIS-reconstituted DRAGA mouse lungs and in a human lung control

Lungs from 10 non-infected HIS-DRAGA and 10 non-infected, non-HIS reconstituted DRAGA mice were homogenized in the presence of MPER mammalian protein extraction reagent (Fisher Scientific, Waltham, MA, USA) containing complete protease inhibitor cocktail tablets (Roche Diagnostics GmbH, Mannheim, Germany) using tubes loaded with ceramic beads (MP Biologicals, Irvine, CA, USA) in a Fast-prep homogenizer (MP Biologicals). Pooled lung homogenates from each group of mice were sonicated on ice in a Fisher Ultrasonicator for 10 cycles of 10 seconds each, the cellular debris was removed by centrifugation at 5,000 rpm, and the protein in the clear supernatant was quantified using a BCA reagent (Thermo Fisher Scientific, Waltham, MA, USA). 9 mg of total lung protein extract from each group of mice and 2 mg of human lung total protein lysate (Zyagen, San Diego, CA, USA) were then individually incubated with gentle shaking (300 strokes per min) in an Eppendorf thermomixer for 1 h at 37°C with 10 μg of the S1(RBD)-mFcγ2a protein (ACRO Biosystems, Newark, DE, USA) followed by incubation with gentle shaking for 1 h at 37°C with 50 μl of rat anti-mouse IgG2a-magnetic microbeads (Miltenyi Biotech, Berdisch Gladback, Germany). The total lysate from each sample was next passed over MACS magnetic columns (Miltenyi Biotec), and the hACE2/S1(RBD)-mFcγ2a/rat anti-mouse IgG2a-magnetic beads were eluted according to the manufacturer’s instructions and concentrated to 52 μl each. The amounts of hACE2 protein in these immunoprecipitates were then quantified using the highly sensitive human ACE2 ELISA kit PicoKine™ (Boster Biological Technology, Pleasanton, CA) per the manufacturer’s protocol. The provided recombinant hACE2 protein was serially diluted in the provided sample buffer. The human lung sample was diluted 1:100 and the DRAGA and HIS-DRAGA samples were diluted 1:50 each in the provided sample buffer. OD450nm values were then read for duplicate samples (100 μl each) using a BioTEK microplate reader (BioTek Instruments, Inc., Winooski, VT, USA). A hACE2 standard curve was constructed by applying a four-parameter logistic fit formula using the BioTEK Gen 5 software (BioTek Instruments, Inc.). The sample OD450nm mean values were then converted to hACE2 concentrations using this curve, per the manufacturer’s instructions.

### Western blot analysis of hACE2 protein in HIS-DRAGA lung lysates

Western blots were next run using aliquots from the same concentrated immunoprecipitates used for the ELISA assays above: immunoprecipitates obtained from (1) 2 mg human lung extract; (2) 9 mg HIS-DRAGA mice lungs lysate; (3) 9 mg DRAGA mice lungs lysate. One μl of each immunoprecipitate added to wells of a 4-12% Bis-Tris gradient pre-cast gel from Invitrogen (Thermo Fisher Scientific, Waltham, MA, USA) and the samples were electrophoresed under denaturing conditions and then electro-transferred onto a PVDF membrane. The membrane was blocked overnight at 4°C with shaking using 5% non-fat milk plus 3% BSA in PBS, incubated with a mouse monoclonal anti-human ACE2 (Abcam, Cambridge, MA, USA, 1:750) for 2h at room temperature, and washed with PBS + 0.01% Tween 20. The membrane was then incubated with goat anti-mouse IgG-HRP (Santa Cruz Biotechnology, Dallas, TX, USA, 1:3000) and SuperSignal™ West Pico PLUS chemiluminescent substrate according to the manufacturer’s instructions (Thermo Fisher Scientific). The chemiluminescent bands were then imaged using a Fluorchem E Imaging System (ProteinSimple, San Jose, CA, USA).

### Infection of mice with SARS CoV-2 virus

HIS-DRAGA mice were infected intranasally (i.n.) with SARS CoV-2 strain USA-WA1/2020 (BEI Resources NR-52281, batch #70033175), courtesy of the Centers for Diseases Control and Prevention (CDC). This virus strain was originally isolated from an oropharyngeal swab of a patient with a respiratory illness who had returned to Washington State, USA, from travel to China and developed COVID-19 in January 2020. Infection of HIS-DRAGA mice and harvesting of serum and organs were conducted in a BSL-3 laboratory at Bioqual, Inc. (Rockville, MD, USA) in compliance with local, state, and federal regulations under IACUC protocol #20-019P. The SARS-COV-2 stock was expanded in Vero E6 cells, and the challenging virus was collected at day 5 of culture when the infection reached 90% cytopathic effect. The full viral genome sequence showed 100% identity with the parent virus sequence listed in GenBank (MN985325.1). A plaque forming assay carried out with confluent layers of Vero E6 cells was used to determine the concentration of live virions, reported as plaque-forming units (pfu). HIS-DRAGA mice were infected i.n. with three different doses (2.8×10^3^, or 2.8×10^4^, or 1×10^3^ pfu) of the same SARS-COV-2 virus strain (NR-52281, batch #70033175) as summarized in **Table S1**.

### Measurement of antibody serum titers to SARS-CoV-2 viral proteins

Titers of human IgM and IgG serum antibodies (1/20 serum dilution) to the recombinant S1(RBD) viral protein from mice infected i.n. with SARS-CoV-2 virions (10^3^ pfu/mouse) were measured prior to infection and at 24 dpi using ELISA kits according to the manufacturer’s instructions (Bethyl Laboratories). In addition, titers of human IgM and IgG serum antibodies in aliquots of these same serum samples against a recombinant His-tagged S trimeric protein and against a recombinant His-tagged N protein (both from ACRO Biosystems) were determined using an in-house ELISA. Briefly, His-S trimeric protein or His-N protein, respectively, were coated on high-binding ELISA plates (Corning Costar, Fisher Scientific, Waltham, MA, USA) at 0.2 μg/well/100μL in carbonate buffer, pH 9.0. The plates were then incubated overnight at 4°C, then blocked with PBS + 1% BSA for 2h at room temperature, washed with PBS + 0.05% Tween 20, and incubated at room temperature for 1h with the serum samples diluted in PBS + 1% BSA + 0.05% Tween 20. Bound human IgM and IgG antibodies against the His-S trimer protein were then revealed by adding anti-human IgM or IgG antibody-HRP conjugates (Bethyl Laboratories) to the His-S-trimer-coated plates. Due to limited serum volumes, total IgG + IgM antibodies against the His-N protein were revealed by adding anti-human kappa plus lambda antibody-HRP conjugates (Bethyl Laboratories) to the His-N protein coated plates. The ELISA plates were then incubated with soluble HRP substrate for 15 minutes, and reactions were stopped by adding H2SO4 (0.18M, 100 μL/well). Plates were read in an ELISA reader at 450nm and 570 nm. OD450nm values were corrected by subtracting the OD570nm values (ranging from 0.045–0.067) of serum samples from the same mice prior to infection. Standard deviations (+/−SD) for each serum sample in duplicate wells were determined at 99% interval of confidence by SigmaPlot v.14 software. The positive control (anti-S1(RBD)) antibody from the kit was used as a positive control for both S1(RBD) and S1-trimer binding ELISA assays. The antibody titer against the N protein in serum from a non-infected mouse served as a negative control for the N-binding ELISA assays.

### Histopathology of lungs from infected HIS-DRAGA mice

Lungs harvested from infected mice at the experimental end-points (14 dpi or 25 dpi) were fixed in 10% formalin, embedded in a paraffin block, and 5 μm sections were stained with Hematoxylin/Eosin (H&E) or Masson’s Trichrome by Histoserv, Inc. (Germantown, MD, USA). Microscopic images were captured using an Olympus BX43 microscope (Shinjuku-ku, Tokyo, Japan).

### Immunofluorescence microscopy

Tissue sections (5μm) from paraffin-embedded cassettes or frozen OCT cassettes from infected and non-infected HIS-DRAGA mice, and from non-infected, non-HIS reconstituted DRAGA mice were prepared by Histoserv, Inc. Thawed OCT-frozen tissue slides were rehydrated with PBS, and paraffin-embedded sections were de-paraffinized with xylene and rehydrated with graded concentrations of ethanol. Slides were then fixed, permeabilized with fixation/permeabilization buffer (Invitrogen, Waltham, MA, USA), blocked with 3% BSA in PBS for 30 min at 37°C, and stained with fluorochrome-conjugated antibodies in PBS containing 0.01% Evans Blue at 37°C for 50 min. To visualize hACE2, slides were probed with the S1(RBD)-mFcγ2a protein (10 μg/ml), washed with PBS, and then incubated with a goat anti-mouse IgG-FITC conjugate (Southern Biotech, Birmingham, AL, USA). Other antibodies to detect antigens of interest were: anti-human CD3-FITC, anti-human CD4-PE, anti-human CD8-PE, anti-human CD45-FITC, anti-human granzyme-B-PE, anti-human CD103-FITC (all from BD Biosciences, San Jose, CA, USA), anti-human CD326-PE (Miltenyi Biotech), anti-human Perforin-PE (Biolegend, San Diego, CA, USA), anti-human CD103-FITC (BD PharMingen, Irvine, CA, USA), anti-mouse CD61-PE (Invitrogen), and anti-hACE2 antibody (clone# MM0073—11A3, Abcam), anti-human TMPRSS2 monoclonal IgG1 mouse antibody (#clone P5H9-A3, Sigma Aldrich), mouse IgG1/κ-Alexa Fluor 594 anti-human CD31 Antibody (clone WM59, Biolegend), goat F(ab’)2 anti-mouse IgG1-PE conjugate (Southern Biotech), and goat F(ab’)2 IgG anti-mouse IgG2a (Southern Biotech). After staining, the slides were washed 3X with PBS, air-dried, and mounted with 10 μl of Vectashield containing DAPI (Vector Laboratories, Burlingame, CA, USA), and images were acquired with a Zeiss Axioscan Confocal microscope or an Olympus BX43 microscope.

---

**Table S1.** Human immune parameters of HIS-DRAGA mice

| Mouse* | Gender | Time post-infusion with human stem cells | Peripheral blood |  | SARS-CoV-2 challenge dose (pfu) | Euthanasia (days post-infection) | Cord blood HLA haplotype<sup>^</sup> |
| --- | --- | --- | --- | --- | --- | --- | --- |
|  |  |  | % human B cells (CD19<sup>+</sup>) | % human T cells (CD3<sup>+</sup>) |  |  |  |
| M#1 | M | 21 weeks | 4.5 | 13.5 | 2.8x10<sup>3</sup> | 1(died) | A |
| F#1 | F | 21 weeks | 48.6 | 9.3 | 2.8x10<sup>3</sup> | 14 | A |
| F#2 | F | 21 weeks | 21 | 33.4 | 2.8x10<sup>4</sup> | 14 | A |
| F#3 | F | 16 weeks | 9.0 | 2.5 | 1x10<sup>3</sup> | 25 | B |
| F#4 | F | 16 weeks | 37.0 | 4.7 | 1x10<sup>3</sup> | 25 | B |
| F#5 | F | 16 weeks | 23.6 | 5.0 | 1x10<sup>3</sup> | 25 | B |
| F#6 | F | 16 weeks | 0.6 | 12.5 | 1x10<sup>3</sup> | 25 | B |
| F#7 | F | 30 weeks | 5.9 | 42.6 | 1x10<sup>3</sup> | 25 | A |
| F#8 | F | 24 weeks | 16.1 | 37.8 | 1x10<sup>3</sup> | 25 | C |
| F#9 | F | 16 weeks | 21.5 | 8.0 | 1x10<sup>3</sup> | 25 | B |
| F#10 | F | 24 weeks | 6.0 | 23.8 | 1x10<sup>3</sup> | 25 | C |
| M#2 | M | 30 weeks | 8.3 | 9.3 | 1x10<sup>3</sup> | 3 | A |
| M#3 | M | 16 weeks | 5.7 | 15.0 | 1x10<sup>3</sup> | 3 | B |
| M#4 | M | 16 weeks | 2.0 | 51.1 | 1x10<sup>3</sup> | 3 | B |
| F#11 | F | 16 weeks | 6.1 | 25.4 | 1x10<sup>3</sup> | 3 | B |
| F#12 | F | 24 weeks | 7.0 | 8.8 | 1x10<sup>3</sup> | 3 | C |
| F#13 | F | 24 weeks | 0.6 | 26.8 | 1x10<sup>3</sup> | 3 | C |
| a | M | 20 weeks | 18.9 | 28.3 | ---- | ---- | D |
| b | M | 20 weeks | 0.9 | 41.4 | ---- | ---- | D |
| c | F | 24 weeks | 52.5 | 17.5 | ---- | ---- | D |
| d | M | 24 weeks | 7.4 | 16.6 | ---- | ---- | D |
| e | M | 24 weeks | 1.0 | 22.1 | ---- | ---- | D |
| f | M | 24 weeks | 1.2 | 44.7 | ---- | ---- | D |
| g | M | 24 weeks | 17.6 | 38.3 | ---- | ---- | D |
| h | F | 20 weeks | 35.2 | 5.0 | ---- | ---- | E |
| i | M | 26 weeks | 4.9 | 26.1 | ---- | ---- | E |
| j | M | 26 weeks | 21.9 | 30.7 | ---- | ---- | E |

Lungs from uninfected mice a-j were pooled to assess human ACE2 mRNA and protein expression.
<sup>^</sup>A: A02:01/A24:02/B13:02/B44:02/DR01:01/DR04:01
<sup>^</sup>B: A01:01/A02:01/B08:01/B15:01/DR03:01/DR04:01
<sup>^</sup>C: A02:01/A24:02/B44:02/B52:01/DR04:01/DR11:04
<sup>^</sup>D: A02:01/A02:01/B18:01/B44:02/DR04:01/DR11:04
<sup>^</sup>E: 02:01/A02:01/B08:01/B27:05/DR03:01/DR04:01

**Figure S1.**
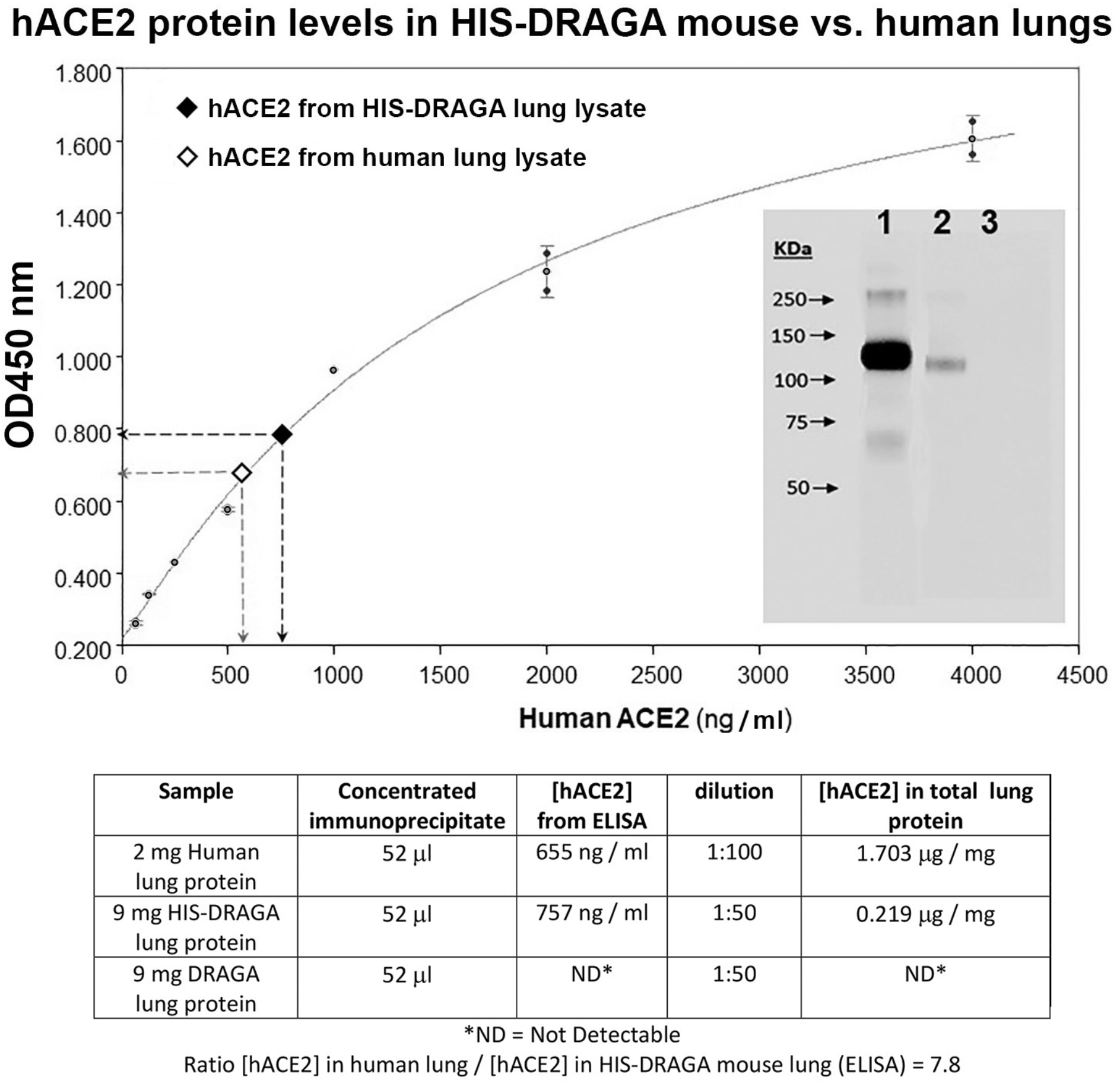
Quantification of hACE2 protein in HIS-DRAGA lungs. Human ACE2 levels in immunoprecipitates obtained from non-infected HIS-DRAGA and human lung lysates using S1(RBD)-mFc◻2a protein + rat anti-mouse IgG2a-magnetic beads were quantified by ELISA. Of note, the OD450nm values for protein immunoprecipitated from a pool of 10 non-infected, non-HIS-humanized DRAGA mouse lung lysates (negative control) fell below the limit of detection (OD450nm <0.05). ***Insert*** shows Western blot detection of hACE2 protein in the concentrated immunoprecipitates probed with a mouse monoclonal anti-human ACE2 IgG followed by goat anti-mouse IgG-HRP with ECL detection. *Lane 1*, human lung immunoprecipitate; *lane 2*, HIS-DRAGA mouse lungs immunoprecipitate; *lane 3*, DRAGA mouse lungs immunoprecipitate (note this sample did not contain detectable hACE2). Lower panel shows the experimental conditions for immunoprecipitation of hACE2, quantification by ELISA, and the ratio of hACE2 in human versus HIS-DRAGA mouse lung samples.

**Figure S2.**
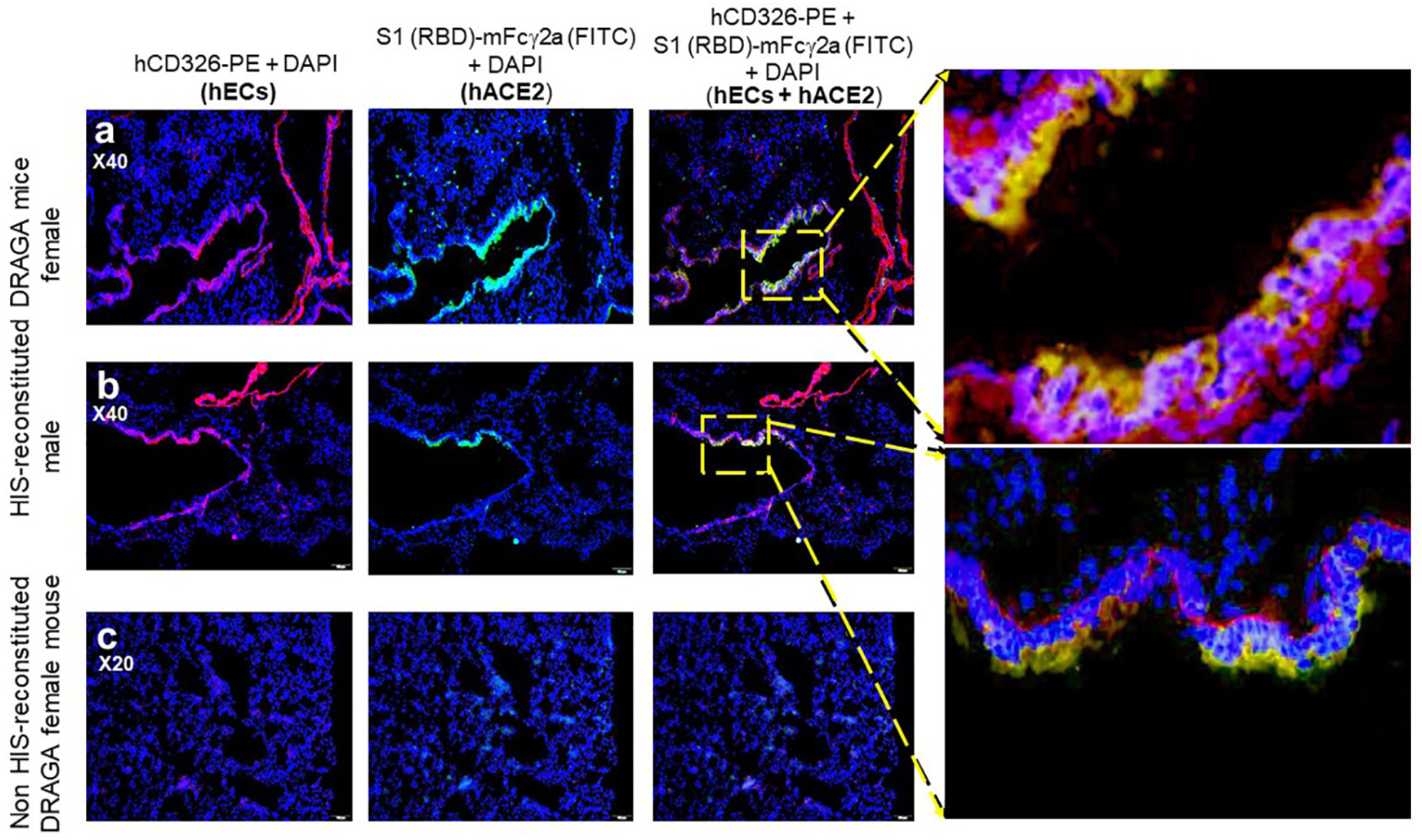
Co-localization of hACE2 receptor with hCD326^+^ alveolar ECs in a non-infected HIS-DRAGA mouse. **a,b.** Co-localization of hACE2 with alveolar hCD326^+^ ECs (orange) revealed by co-staining of S1(RBD)-mFc◻2a protein + goat anti-mouse IgG-FITC (green) and anti-hCD326-PE (red) in representative HIS-DRAGA female and male mice. **c.** Lack of hCD326^+^ ECs and negligible binding of S1(RBD)-mFc◻2a protein to a representative lung section from a non-HIS-reconstituted DRAGA female mouse.

**Figure S3.**
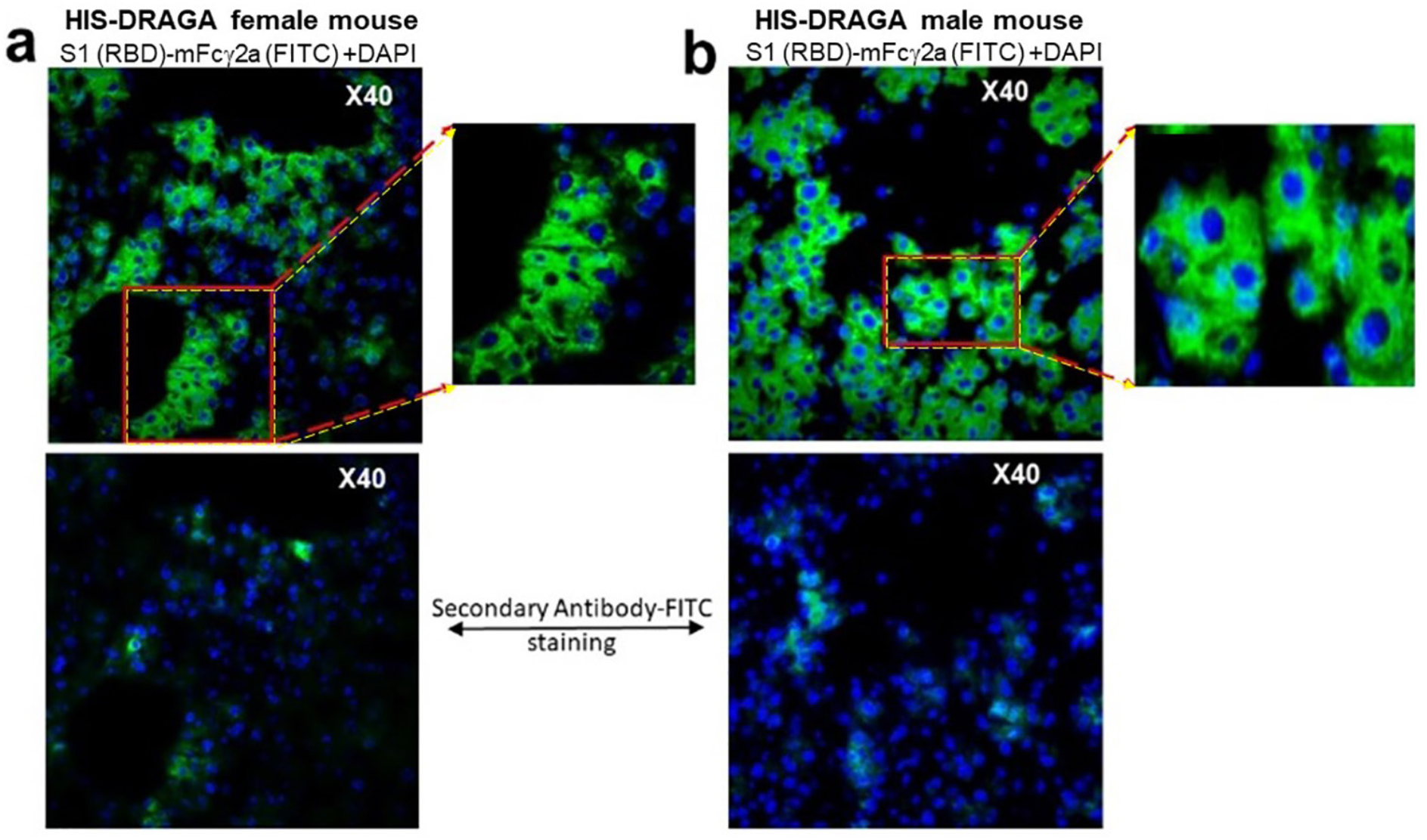
Binding of SARS-CoV-2 S1(RBD) protein to the endothelium of liver cholangiocytes in non-infected HIS-DRAGA mice. S1(RBD) binding to the liver cholangiocytes from representative non-infected HIS-DRAGA female (*panel a)* and male *(panel b)* mice. Merged images and enlargements show binding of S1(RBD)-mFc◻2a revealed by a goat anti-mouse IgG-FITC conjugate (green) and nuclei (DAPI, blue). *Lower* panels, representative images showing minimal background binding of the goat anti-mouse IgG-FITC secondary antibody (green) and nuclei (DAPI, blue) in tissues from the same mice in panels a and b.

**Figure S4.**
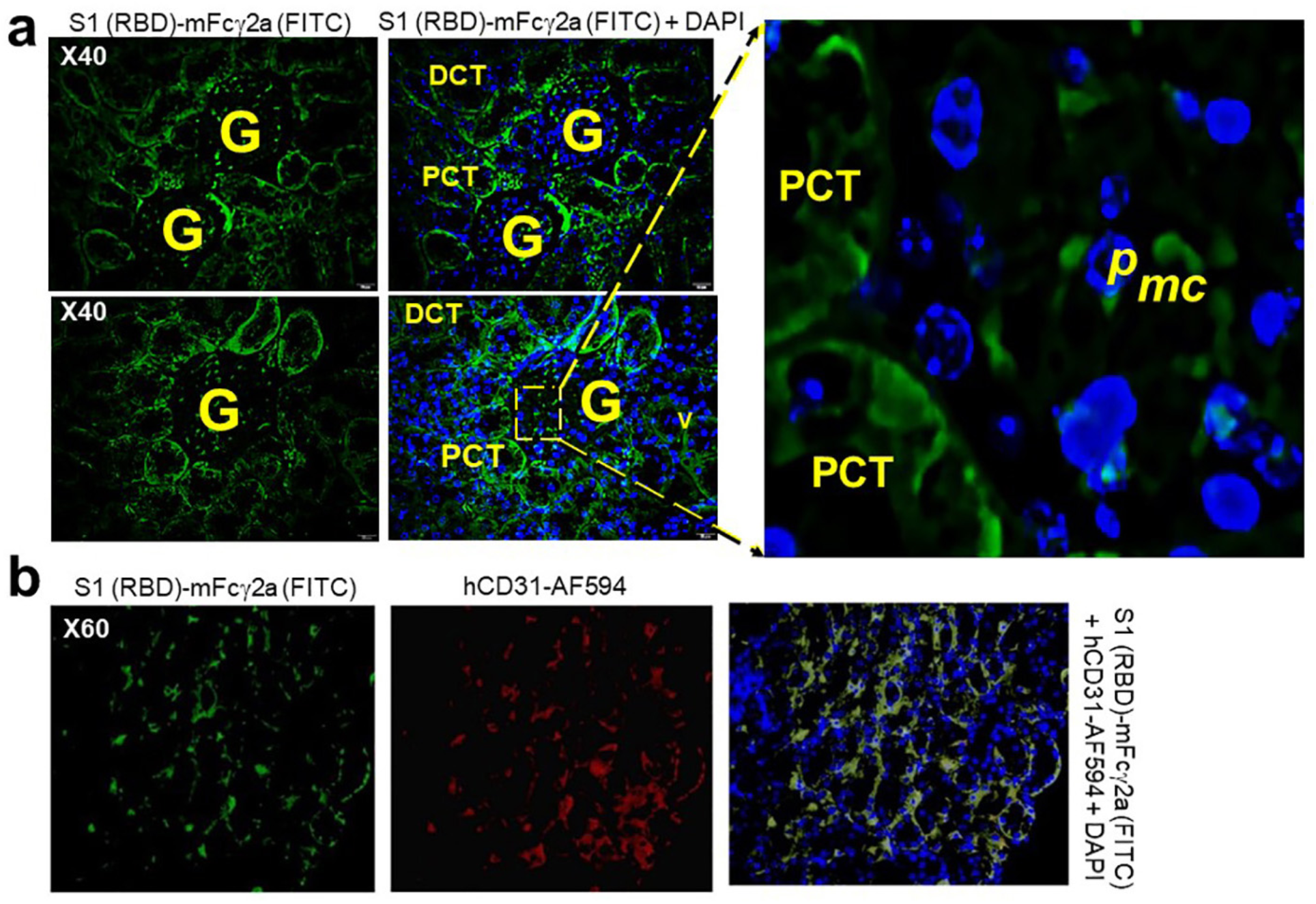
Binding of SARS-CoV-2 S1 (RBD) protein to kidney epi/endothelia of infected HIS-DRAGA mice. **a.** Sections of renal cortex from HIS-DRAGA survivors #F1 (upper panels) and #F2 (lower panels) of SARS-CoV-2 infection with 2.8×10^4^ pfu and 2.8×10^3^ pfu, respectively, at the experimental endpoint (14 dpi). Sections were co-stained with DAPI (blue) and S1(RBD)-mFc◻2a protein + goat anti-mouse IgG-FITC (green). Shown is the S1(RBD) protein bound to the epithelium layer of proximal convoluted tubules (PCT) and distal convoluted tubules (DCT) surounding the glomeruli (G). Enlargement of a peripheral glomerular area shows nuclei (blue) of podocytes (p) and the endothelia (green) of glomerular microcapilaries (mc) in close proximity to the podocytes (green). **b.** Expression of hACE2 revealed by S1(RBD)-mFc◻2a binding (green) on kidney epithelial cells labeled with an anti-hCD31-AF594 antibody (red).

**Figure S5.**
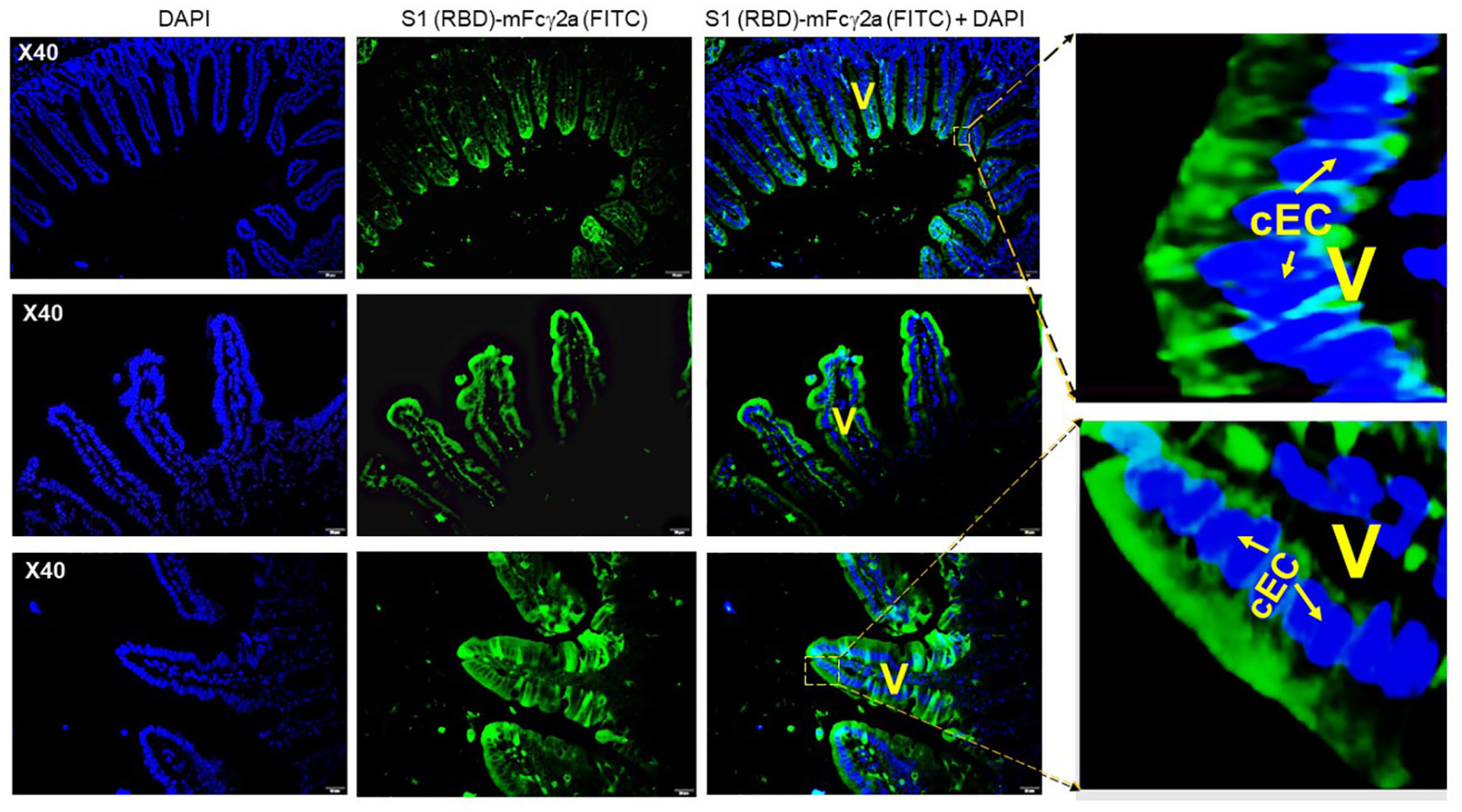
Binding of SARS-CoV-2 S1(RBD) protein to intestinal epithelia of infected HIS-DRAGA mice. Tissue sections from the small intestine of HIS-DRAGA mouse #F1 (*upper and middle panels*) and #F2 (*lower panel*) survivors of SARS-CoV-2 infection with 2.8×10^4^ pfu and 2.8×10^3^ pfu, respectively, at the experimental endpoint (14 dpi). Sections were co-stained with DAPI (nuclei, blue) and S1(RBD)-mFc◻2a protein+ goat anti-mouse IgG-FITC (green). Shown is the S1(RBD) protein bound to the columnar epithelial cells (cEC) of the absorptive intestinal villi (V).

**Figure S6.**
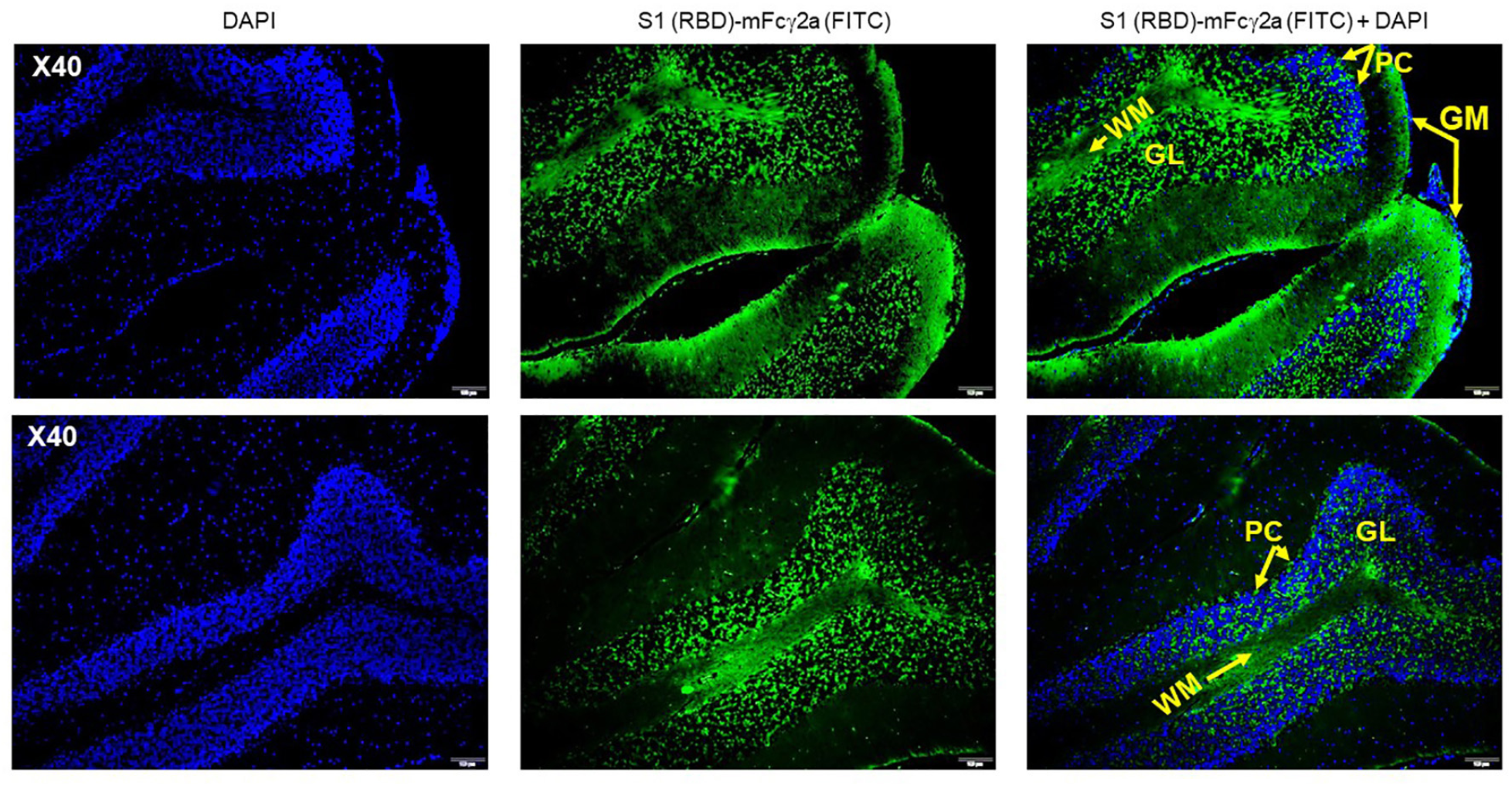
Binding of SARS-CoV-2 S1(RBD) protein to the brain epithelia in infected HIS-DRAGA mice. Tissue sections of cerebellum cortex from HIS-DRAGA mice #F1 (*upper panel*) and #F2 (*lower panel*) survivors of SARS-CoV-2 infection with 2.8×10^4^ pfu and 2.8×10^3^ pfu, respectively, at the experimental endpoint (14 dpi) co-stained with DAPI (nuclei, blue) and S1(RBD)-mFc◻2a protein + goat anti-mouse IgG-FITC (green). Shown is the S1(RBD) protein bound to the white matter (WM), granular layer (GL), and Purkinje cells (PC), but not to the outer neuronal layer in the grey matter (GM).

**Figure S7.**
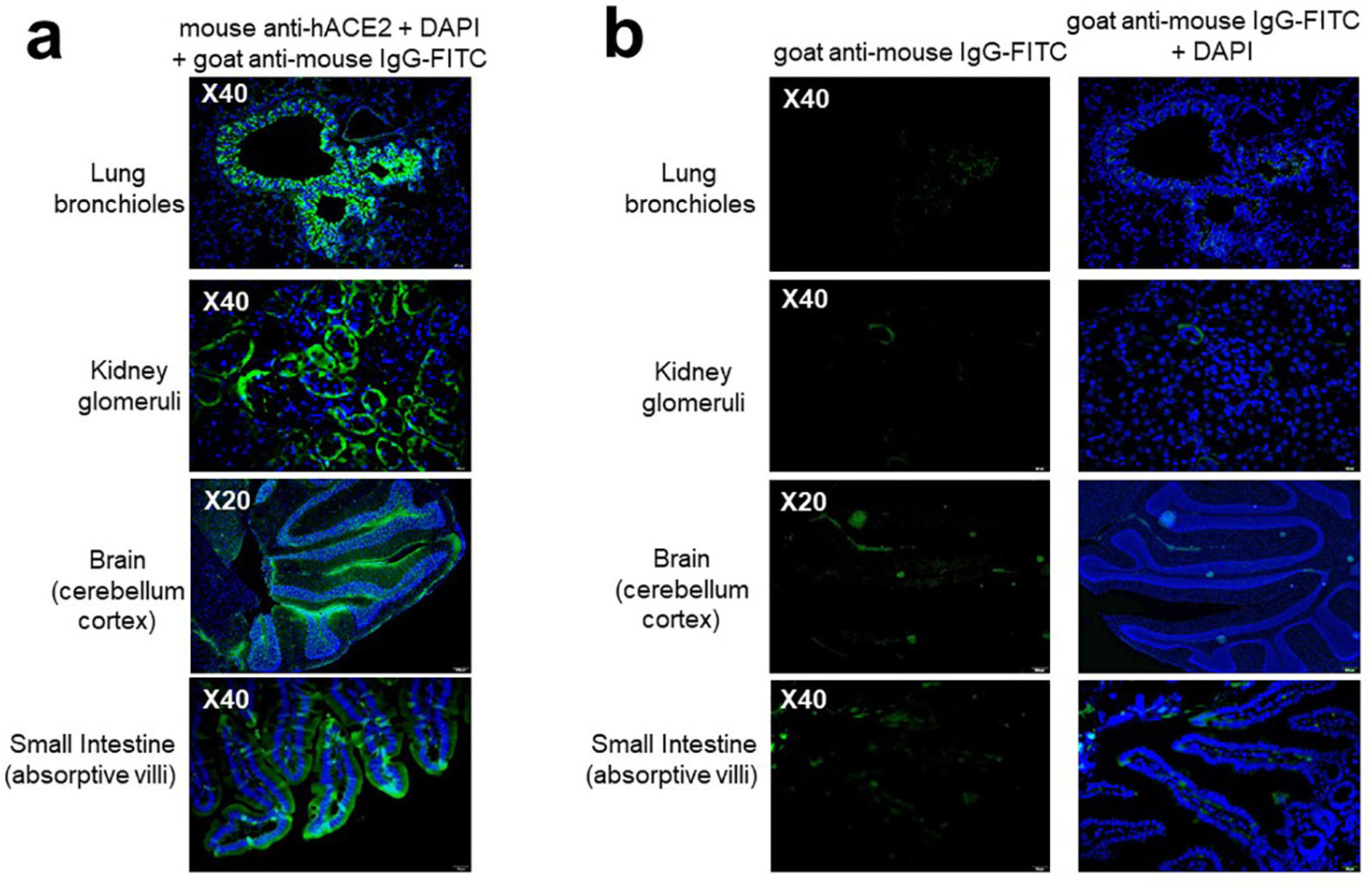
Identification of hACE2 on tissue sections from organs of a non-infected HIS-DRAGA mouse, detected with an anti-hACE2 specific antibody. **a.** Merged images of tissue sections from a representative non-infected HIS-DRAGA mouse stained with mouse anti-hACE2 followed by goat anti-mouse IgG-FITC (green) and DAPI (blue). **b.** Minimal background binding of the secondary antibody (goat anti-mouse IgG-FITC) (left) overlapped with DAPI staining (right) of tissue sections adjacent to those shown in *panel a*.

**Figure S8.**
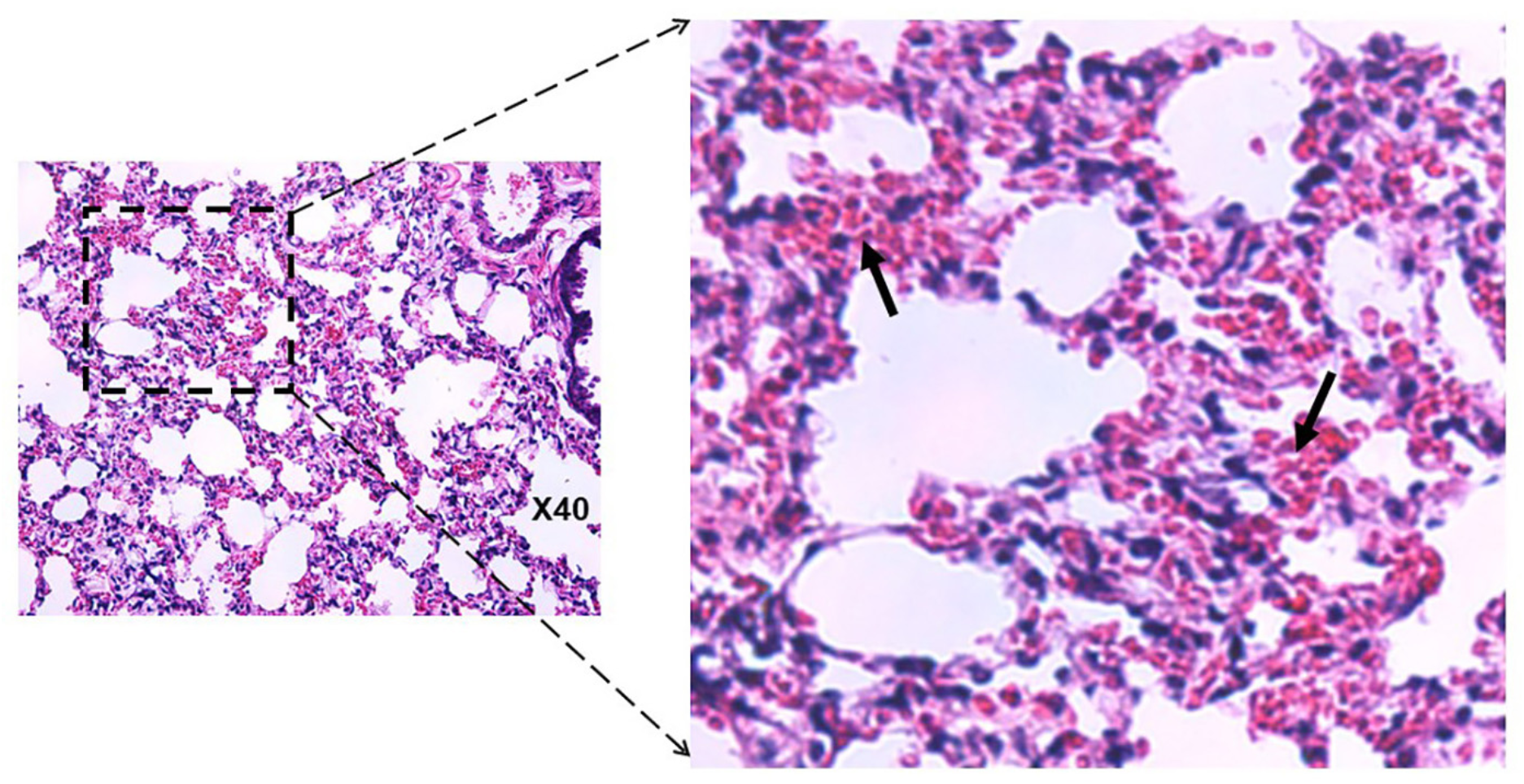
Lung pathology of a HIS-DRAGA mouse recovering from SARS-CoV-2 infection. Representative H&E-stained lung section from HIS-DRAGA mouse #F2 that recovered its initial body weight at 9 days after infection with SARS-CoV-2 at 2.8×10^3^ pfu. Interstitial and intra-alveolar infiltrates are indicated by arrows.

**Figure S9.**
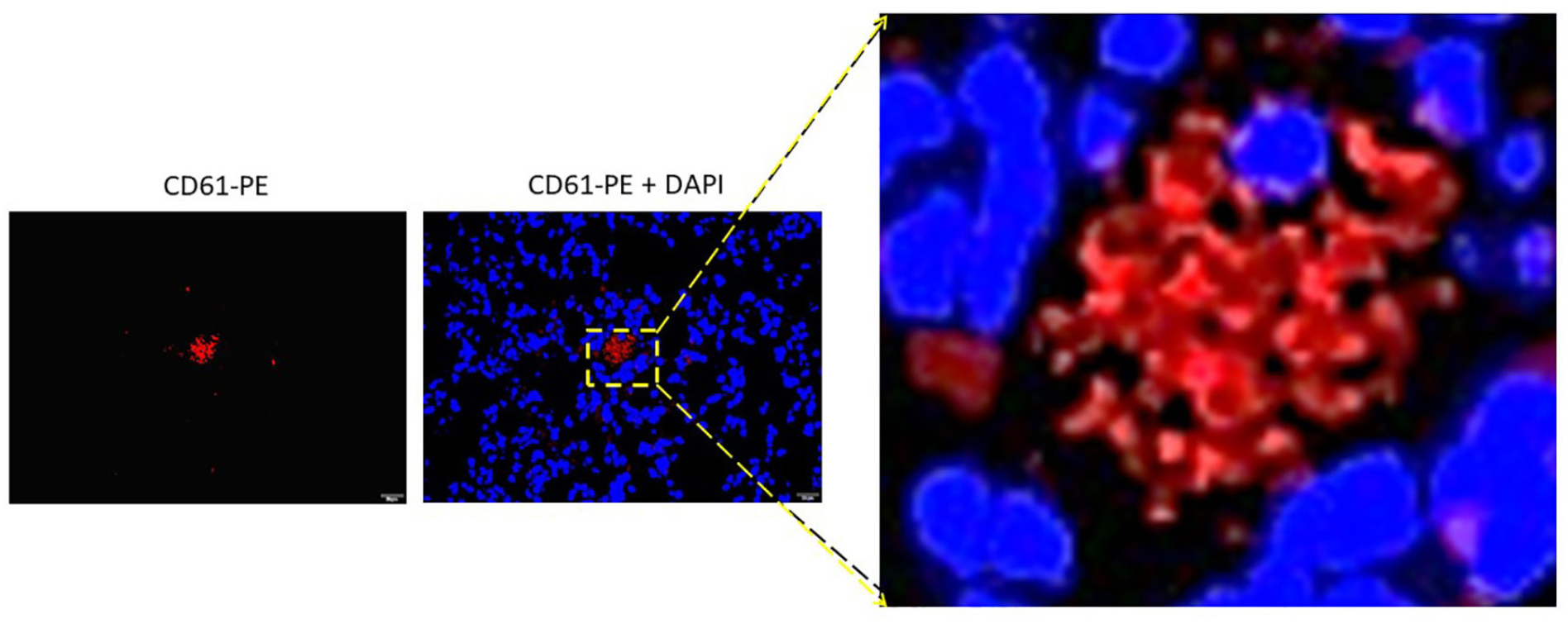
CD61^+^ intra-alveolar microthrombi in a SARS-CoV-2 infected HIS DRAGA mouse. Intra-alveolar microthrombi in lung section from a HIS-DRAGA mouse infected with 2.8×10^4^ pfu of SARS-CoV-2, which had not recovered its initial body weight by 14 dpi. *Left panel*, staining with anti-CD61-PE (red). *Right panel* with enlargement: merged anti-CD61-PE co-staining with DAPI (blue). The non-nucleated CD61^+^ cluster indicates this is a platelet micro-thrombus.

**Figure S10.**
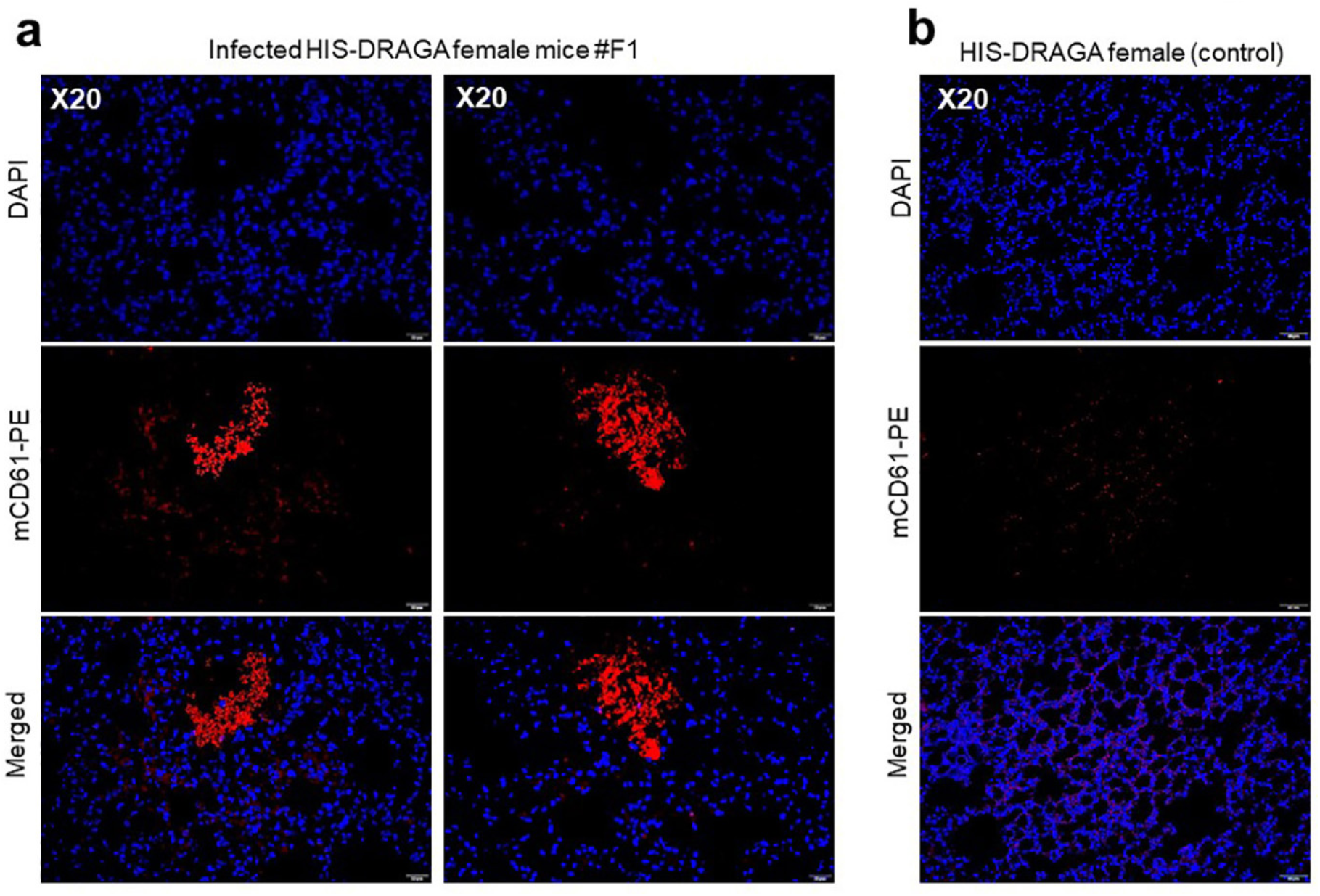
Large intra-alveolar thrombi in a SARS-CoV-2 infected HIS-DRAGA mouse. **a.** Representative lung section from HIS-DRAGA mouse #F1 infected with 2.8×10^4^ pfu, which had not recovered its initial body weight by 14 dpi. Co-staining with anti-CD61-PE (red) + DAPI (blue). **b.** Representative lung section from a non-infected HIS-DRAGA mouse stained as in panel a, with no evidence of intra-alveolar thrombi.

**Figure S11.**
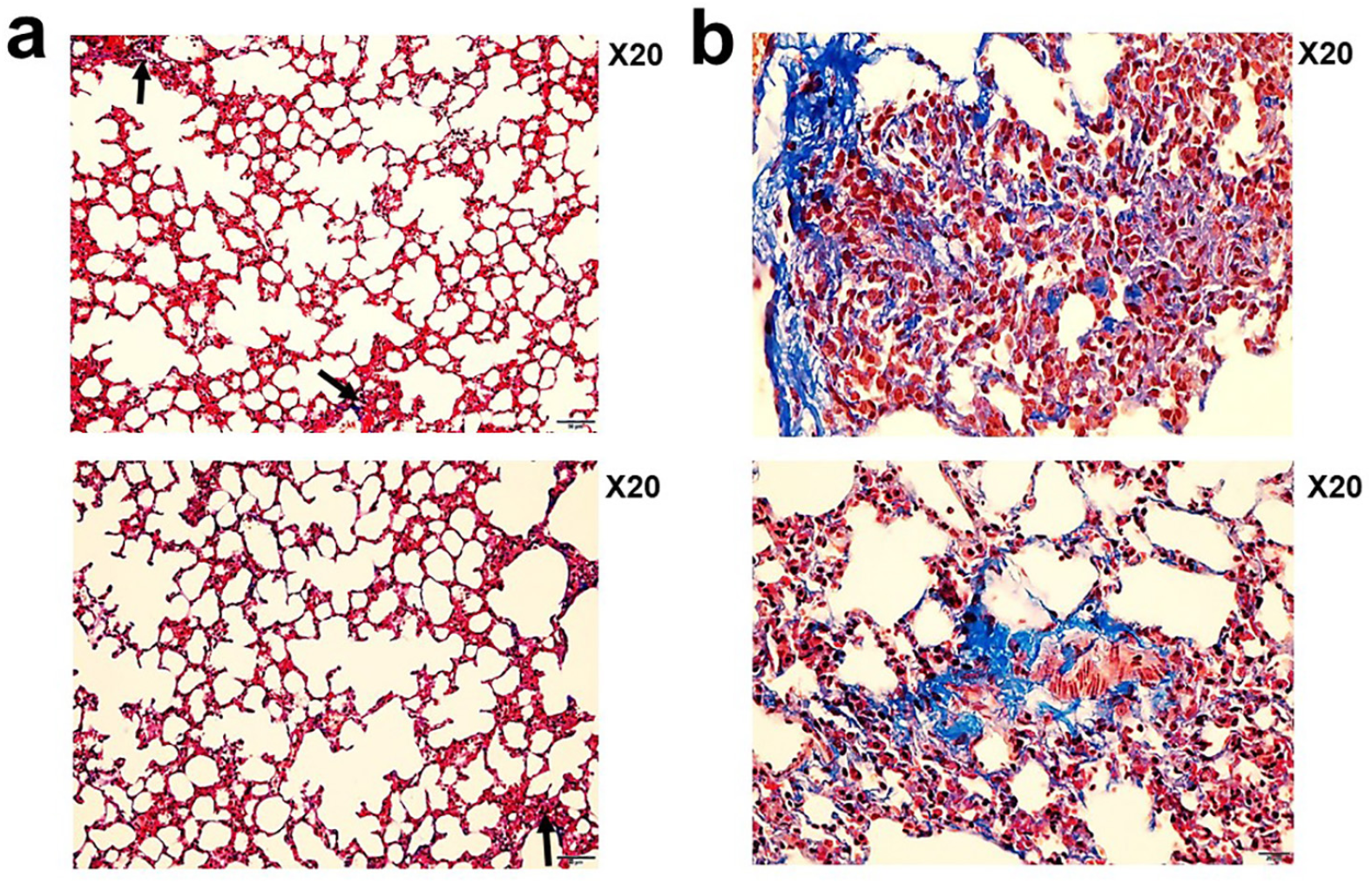
Pulmonary sequelae in SARS-CoV-2 infected DRAGA mice. **a.** Masson’s Trichrome staining of lung sections from HIS-DRAGA mice #F3 (upper panel) and #F4 (lower panel) infected with 10^3^ pfu SARS-CoV-2, which recovered their initial body weights by 9 and 25 dpi, respectively. Shown are small peri-alveolar infiltrates building collagen fibers (blue, arrows). **b.** Masson’s Trichrome staining of lung sections from HIS-DRAGA mice #F5 (*upper panel*) and #F6 (*lower panel*) infected with 10^3^ pfu SARS-CoV-2, which had not recovered their initial body weights by 25 dpi. Shown are peri-alveolar and intra-alveolar infiltrated areas building collagen fibers (blue).

**Figure S12.**
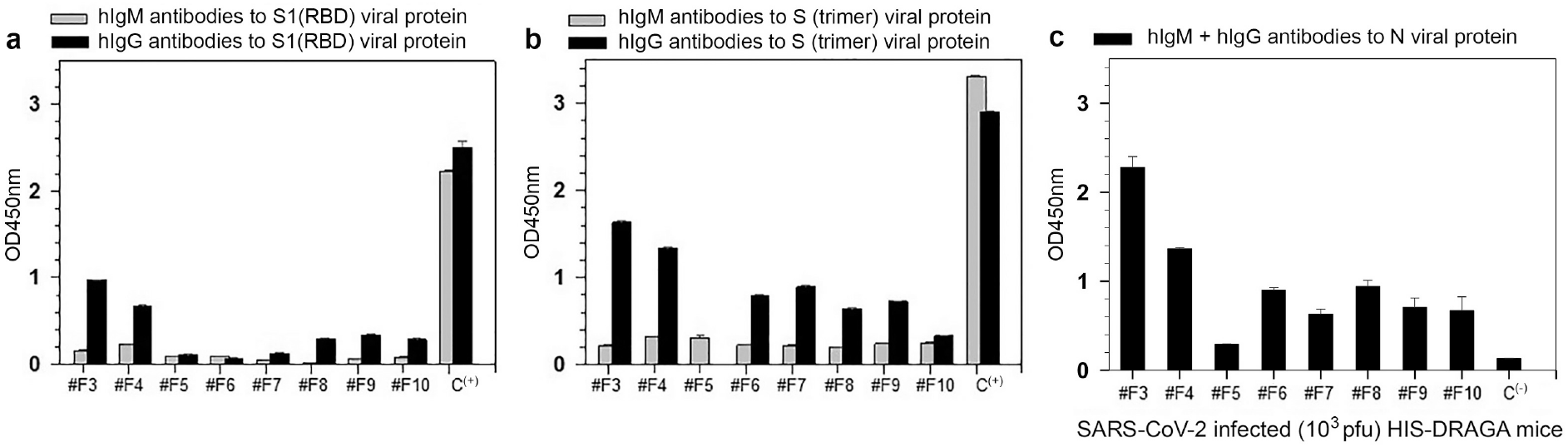
Antibody serum titers to SARS-CoV-2 viral proteins. Titers of hIgM and hIgG antibodies to S1(RBD) protein (**a**) S-trimer protein (***b***) and N protein (**c)** in sera (diluted 1:20) from 8 HIS-DRAGA mice infected with SARS-CoV-2 (10^3^ pfu), as measured by ELISA at 25 dpi. An anti-S1(RBD) antibody provided in the kit (Bethyl Laboratories) was included as a positive control in panels a and b (C+). The antibody titers against the N protein in serum from a non-infected mouse served as a negative control in panel c. OD450nm values were corrected by subtracting values (ranging from 0.045–0.067) of serum samples from the same mice prior to infection. Standard deviations (+/−SD) for each serum sample in duplicate wells were determined at 99% interval of confidence by SigmaPlot v.14 software.

KEY REAGENTS AND RESOURCES

KEY REAGENTS AND RESOURCES
| Kits, Recombinant proteins and Chemicals | Source | Catalog # |
| --- | --- | --- |
| EasySep Human CD3 Positive Selection Kit | Stem Cell Technologies, Vancouver, BC, Canada | 18051 |
| Qiagen RNA Extraction Kit | Qiagen, Hilden, Germany | 74104 |
| Human lung mRNA | Thermo Fisher Scientific, Waltham, MA, USA | AM7968 |
| One-Step RT-PCR kit | Qiagen, Hilden, Germany | 210210 |
| MPER mammalian protein extraction reagent | Thermo Fisher Scientific, Waltham, MA, USA | 78501 |
| Complete protease inhibitor cocktail tablets | Roche Diagnostics GmbH, Mannheim, Germany | 1186153001 |
| Ceramic beads | MP Biologicals, Irvine, CA, USA | 6913100 |
| Fast-prep homogenizer | MP Biologicals, Irvine, CA, USA | SKU 116005500 |
| BCA reagent | Thermo Fisher Scientific, Waltham, MA, USA | 23225 |
| S1(RBD)-mFcgamma2a chimeric antibody | ACRO Biosystems, Newark, DE, USA | S1N-C5257 |
| Rat anti-mouse IgG2a-magnetic microbeads | Miltenyi Biotec, Berdich Gladback, Germany | 130-047-202 |
| MACS magnetic columns | Miltenyi Biotec, Berdich Gladback, Germany | 130-042-401 |
| SARS-CoV-2 S protein, His Tag, Super stable trimer | ACRO Biosystems, Newark, DE, USA | SPN-C52H9 |
| SARS-CoV-2 N protein, His Tag | ACRO Biosystems, Newark, DE, USA | NUN-C51H9 |
| Bovine serum albumin (BSA) | Sigma-Aldrich, St. Louis, MO, USA | A7030-100g |
| SuperSignal™ West Pico PLUS Chemiluminescent Substrate | Thermo Fisher Scientific, Waltham, MA, USA | 34579 |
| NuPAGE™ 4 to 12%, Bis-Tris, 1.5 mm, Mini Protein Gel, 10-well | Thermo Fisher Scientific, Waltham, MA, USA | NP0335PK2 |
| Human lung total protein | Zyagen, San Diego, CA, USA | HT-601 |
| iBlot™ 2 Gel Transfer Device | Thermo Fisher Scientific, Waltham, MA, USA | IB21001 |
| Vectashield containing DAPI | Vector Laboratories, Burlingame, CA, USA | H-1200-10 |
| Tissue fixation/permeabilization buffer | Thermo Fisher Scientific, Waltham, MA, USA | 00-8333-56 |
| microplate reader | BioTek Instruments, Winooski, VT, USA |  |
| Fluorchem E imaging system | ProteinSimple, San Jose, CA, USA |  |
| RNA-STAT-60 extraction reagent | Tel-Test, Inv., Friendswood, TX, USA | TL-4200 |
| RT-qPCR primers + probe for SARS-CoV-2 Nucleocapsid RNA | Integrated DNA Technologies, Coralville, IA, USA | none |
| RT-qPCR kit | Meridian Bioscience, Memphis, TN, USA | BIO-78005 |

| Antibodies | Source | Catalog # |
| --- | --- | --- |
| Mouse monoclonal antibody to hACE2 | Abcam, Cambridge, MA, USA | Ab-89111 |
| Mouse anti-human CD3-FITC | BD Biosciences, SanJose, CA, USA | 555339 |
| Mouse anti-human CD45-FITC | BD Biosciences, SanJose, CA, USA | 347463 |
| Mouse anti-human CD8-PE | BD Biosciences, SanJose, CA, USA | 555635 |
| Mouse anti-human granzyme-B-PE | BD Biosciences, SanJose, CA, USA | 561142 |
| Recombinant human anti-human CD326-PE | Miltenyi Biotec, Berdisch Gladback, Germany | 130-110-999 |
| Mouse anti-human Perforin-PE | Biolegend, San Diego, CA, USA | 353304 |
| Mouse anti-human CD103-FITC | BD Pharmingen, Irvine, CA, USA | 550259 |
| Hamster anti-mouse CD61-PE | Thermo Fisher Scientific, Waltham, MA, USA | 12-0611-82 |
| Mouse anti-human CD31- Alexa Fluor 594 | Biolegend, San Diego, CA, USA | 303126 |
| Anti-human TMPRSS2 Antibody | Sigma-Aldrich, St. Louis, MO, USA | MABF2158-100UG |
| Goat anti-Mouse IgG Fc-FITC | Southern Biotech, Birmingham, AL, USA | 1013-02 |
| Goat anti mouse IgG-HRP | Santa Cruz Biotechnology, Dallas, TX, USA | Sc-2005 |
| Goat F(ab')<sub>2</sub> anti-mouse IgG2a, Human ads-FITC | Southern Biotech, Birmingham, AL, USA | 1082-02 |
| Goat F(ab')<sub>2</sub> anti-mouse IgG1, Human ads-PE | Southern Biotech, Birmingham, AL, USA | 1072-09 |
| Anti-human IgM antibody-HRP | Bethyl Laboratories, Montgomery, TX, USA | A80-100P |
| Anti-human IgG antibody-HRP | Bethyl Laboratories, Montgomery, TX, USA | A80-104P |
| Anti-human kappa antibody-HRP | Bethyl Laboratories, Montgomery, TX, USA | A80-115P |
| Anti-human lambda antibody-HRP | Bethyl Laboratories, Montgomery, TX, USA | A80-116P |

| ELISA Kits and supplies | Source | Catalog # |
| --- | --- | --- |
| Human ACE2 ELISA kit PicoKine | Boster Biological Technology, Pleasanton, CA, USA | EK0997 |
| SARS-CoV-2 IgM ELISA Kit (antigen is rS1(RBD)) | Bethyl Laboratories, Montgomery, TX, USA | E88-302 |
| SARS-CoV-2 IgG ELISA Kit (antigen is rS1(RBD)) | Bethyl Laboratories, Montgomery, TX, USA | E88-301 |
| Coming Costar high-binding ELISA plates | Thermo Fisher Scientific, Waltham, MA, USA | 07-200-721 |

| Biological samples |  |  |
| --- | --- | --- |
| Umbilical cord units (HLA-A2.1.HLA-DR0401) | New York Blood Center, Long Island City, NY, USA | n/a |

| DRAGA mice and SARS-CoV-2 virus |  |  |
| --- | --- | --- |
| DRAGA mice | WRAIR/NMRC, Silver Spring, MD, USA | n/a |
| SARS-CoV-2 strain USA-WA1/2020 | via U.S. Centers for Disease Control and Prevention, BEI Resources NR-52281 | batch #70033175 |

| Services |  |
| --- | --- |
| Sequencing PCR amplicons | Eurofins, Inc., Coralville, IA, USA |
| Tissue sections from paraffin-embedded organs | Histoserv Inc., Germantown, MD, USA |
| H&E Staining | Histoserv Inc., Germantown, MD, USA |
| Masson's Trichrome staining | Histoserv Inc., Germantown, MD, USA |

